# Key determinants of T cell epitope recognition revealed by TCR specificity profiles

**DOI:** 10.1101/2025.11.17.688817

**Authors:** Yan Liu, Giancarlo Croce, Daniel Tadros, Dana Moreno, Alexandra Michel, Anne-Christine Thierry, Raphael Genolet, Marta AS Perez, Rachid Lani, Philippe Guillaume, Michael Hebeisen, Matei Teleman, Kelvin Lau, Amede Larabi, Julien Racle, Daniel E Speiser, Florence Pojer, Steve Dunn, Petra Baumgaertner, Vincent Zoete, Alexandre Harari, David Gfeller

## Abstract

Interactions between T-Cell Receptors (TCRs) and antigenic peptides presented on Major Histocompatibility Complex (MHC) molecules are central to the immune recognition of infected and malignant cells. The complexity of the TCR sequence space, the flexibility of the TCR-epitope interface and the lack of a standardized framework to visualize TCR specificity have hindered a comprehensive understanding of the principles governing TCR-epitope recognition across a broad range of epitopes. Here, we introduce a fully interpretable probabilistic framework, termed TCR specificity profiles (TSPs), which captures fundamental properties distinguishing epitope-specific from baseline TCR repertoires. We demonstrate that TSPs unravel key determinants of TCR-epitope recognition specificity. By identifying and analyzing TCRs recognizing dozens of epitope variants, we show that TSPs accurately predict cross-reactivity and reveal how TCR specificity evolves with epitope sequence, binding mode and MHC restriction. TSPs further enable interpretation of machine learning tools and reveal how AlphaFold3 can be used to decipher key determinants of TCR-epitope recognition specificity.

## Introduction

T-cell receptors (TCRs) recognize epitopes which consist of antigenic peptides displayed on Major Histocompatibility Complex (MHC) molecules. This recognition is essential for specific cellular immune responses against infections and cancer. To cope with highly diverse epitopes, T cells have the ability to generate heterodimeric TCRs with different sequences through a process called V(D)J recombination. First, specific V and J genes are selected from a pool of germline encoded segments and assembled into a single reading frame for both the alpha and the beta chain (with an additional small D segment in TCRβ chains). The theoretical diversity of Vα/Jα/Vβ/Jβ combinations is in the order of 10^6^. Second, insertions and deletions occur at the V(D)J junction which is comprised within the Complementary Determining Region 3 (CDR3) loop in each chain, thereby further increasing the CDR3 length and sequence diversity.

TCRs recognize epitopes through several loops, called CDR1, CDR2 and CDR3, in both chains. Structural analyses have suggested that residues in CDR1 and CDR2 loops (which are fully encoded in V usage) preferentially make contacts with the MHC, and residues in CDR3 loops preferentially interact with the peptide, although many counter-examples have been documented ^1^. This has led some studies to hypothesize that V usage would primarily encode MHC recognition specificity and CDR3 residues would encode peptide recognition specificity ^2–4^. CDR loops show high flexibility. This flexibility has represented an important limitation to structurally model TCR-epitope complexes at high enough resolution to infer specificity.

Analysis of several epitope-specific TCR sequence datasets has revealed quantifiable features characterizing TCRs recognizing distinct epitopes ^5,6^. Many studies exploring such data have focused on CDR3 sequences, using analytical tools such as binding motifs, k-mers, convolutional neural networks or other machine learning approaches ^6–9^. Diverse statistical patterns have emerged from these analyses and some attempts have been proposed to visualize them ^5,10^. However, standardized frameworks to represent these patterns in a consistent and interpretable way throughout multiple epitopes are still missing.

Epitope-specific TCRs have been used to train many sequence-based machine learning TCR-epitope recognition predictors ^8,11–13^. These tools have reached reasonable accuracy for several epitopes with abundant training data, but predictions remain unreliable for epitopes with little or no training data ^14,15^. Application of powerful AI models for protein structure and interaction predictions, such as AlphaFold (AF) ^16^, shows convincing promises even for epitopes without experimentally known TCRs ^17–23^. Unfortunately, interpretation of both sequence-based and structure-based TCR-epitope interaction predictions remains challenging. This has limited our ability to investigate the most important determinants of TCR-epitope recognition specificity across a broad range of epitopes.

Here, we introduce a probabilistic framework to visualize and understand fundamental properties distinguishing epitope-specific TCRs from large datasets of TCRs with unknown specificities (referred to as ‘baseline TCR repertoires’). We demonstrate that this framework captures key determinants of TCR-epitope recognition specificity and cross-reactivity and enables interpretation of machine learning and AF3-based predictions.

## Results

### TCR specificity profiles accurately capture specificity in T cell epitope recognition

When modelling specificity in epitope recognition by native TCRs, a key step is to find patterns that distinguish epitope-specific TCRs from baseline TCR repertoires. This comparison is especially relevant because baseline TCR repertoires exhibit high diversity in many of their properties. For instance, some V or J segments occur at high (>10%) frequency in TCR repertoires, while others occur at very low (<0.1%) frequency (Figure S1A). In addition, biases related to sequencing protocols, sample processing, patient genotype and patient immune history impact properties of baseline TCR repertoires and lead to variability in V and J usage ^24^.

To systematically address these different issues, we collected hundreds of baseline TCR repertoires from many different samples, representing more than 10^6^ alpha chains and more than 10^8^ beta chains in human and mouse (see Materials and Methods and Table S1). We derived mean (hereafter denoted with the letter 𝑄) and standard deviation for V usage, J usage, CDR3 length distribution and CDR3 amino acid frequencies at each position for each chain. As both CDR3 lengths and amino acid sequences are influenced by V and J usage, we further derived residual probabilities for the CDR3 length distributions (i.e., 𝑄(𝐿|𝑉, 𝐽)) and CDR3 amino acid frequencies (i.e., 𝑄(𝐶𝐷𝑅3|𝑉, 𝐽, 𝐿)) (for clarity the chain and species information is omitted in the formula).

We then developed a fully interpretable visualization framework, termed TCR specificity profiles (TSPs), to compare properties of epitope-specific TCRs with those of baseline TCR repertoires (Figure 1A). First, V and J usage in epitope-specific TCRs are compared to the mean and standard deviation of V and J usage in baseline TCR repertoires. Second, CDR3 length distributions in epitope-specific TCRs are compared to the expected distributions in baseline repertoires with the same V/J usage (see Materials and Methods). Third, CDR3 amino acid frequencies are compared to those in baseline repertoires with the same V/J usage and CDR3 length using sequence motifs (Figure 1A for the dominant CDR3 length, see Figure S1B for other lengths). The last two comparisons ensure that any specificity signal in CDR3 lengths or sequences is not confounded by specificity in V/J choices. Using the example of TCRs recognizing a Yellow Fever epitope (LLWNGPMAV restricted HLA-A*02:01, hereafter referred to as YF) (Figure 1A), one can immediately appreciate highly specific patterns emerging in V/J usage and CDR3 (especially CDR3α) length distributions. Some of these patterns (e.g., enrichment in TRAV12-2 or in CDR3α of length 10) are consistent with previous observations ^25^, while others had not been reported, mainly because of the difficulty to detect enrichment in epitope-specific TCRs without a robust model of baseline TCR repertoires. Similar observations can be made for other epitopes, many of which show strong specificity in some V or J usage that cannot be explained by variability in baseline repertoires (see Figure S1C and https://tcrmotifatlas.unil.ch).

**Figure 1:**
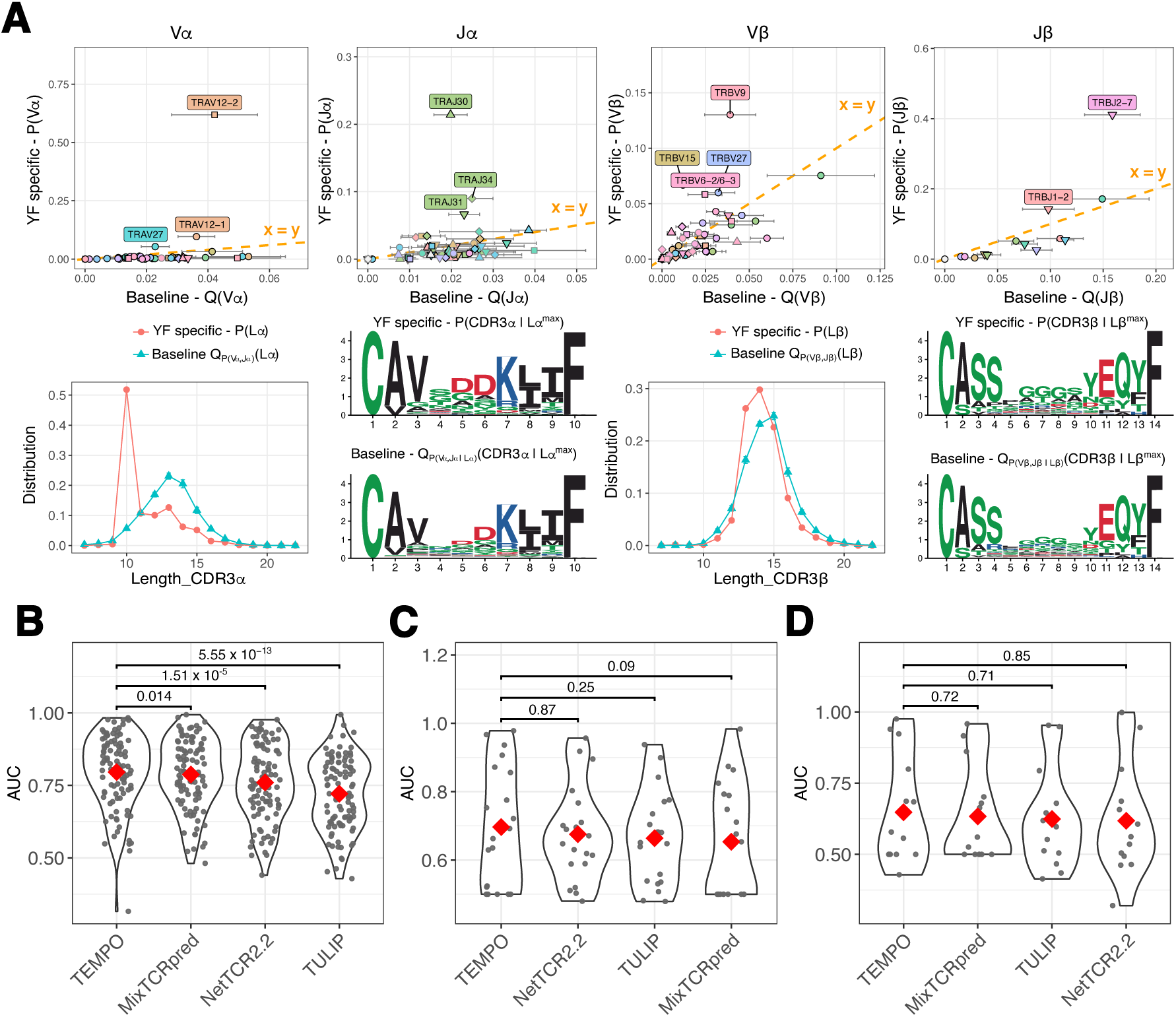
TSPs accurately capture specificity in T cell epitope recognition. **A**: Illustration of the TSP visualization framework based on TCRs recognizing the YF epitope. **B**: AUC for the cross-validation analysis with TEMPO and other tools retrained on the same data. Each point represents an epitope with at least 20 paired TCRs. **C**: AUC obtained with TEMPO and the published version of other tools on the IMMREP23 dataset. **D**: AUC obtained with TEMPO and the published version of other tools on the ePytope-TCR dataset. The red points show the mean values, which have been used to order the tools. P-values represent Wilcoxon paired tests, which are influenced both by the differences in AUC values and by the number of epitopes considered in each benchmark.

To explore how accurately TSPs capture specificity in TCR-epitope recognition, we developed a TCR-epitope interaction predictor where every coefficient can be traced back to the TSPs. This model, called TEMPO for <u>T</u>CR-<u>e</u>pitope <u>m</u>otif-based interaction <u>p</u>redict<u>o</u>r, is derived from the analytical solution of the log-likelihood maximization used in well-established binding motifs. In our case, the different features (i.e., V, J, CDR3 length, CDR3 amino acids) have different alphabets and renormalization is performed with the baseline distributions used in the TSPs for each epitope (see Materials and Methods).

To compare the predictive power of TEMPO with other tools, we compiled a dataset of nearly 100’000 TCRs recognizing hundreds of epitopes (see Materials and Methods and Table S2). For each epitope with at least 20 complete paired TCRαβ sequences, we split these TCRs into five groups and iteratively used four groups for training and one group for testing. To avoid leakage between training and testing data, any TCR with the same CDR3α or same CDR3β as a TCR in the test set was excluded from the training set (see Materials and Methods). Negatives in the test set were further selected from baseline TCR repertoire studies not included in the derivation of the baseline model used in TEMPO (see Materials and Methods). Training and testing on these data both TEMPO and other state-of-the-art machine learning tools ^8,11^, including our own deep learning MixTCRpred ^12^, revealed similar or higher Area Under the Curve (AUC) for TEMPO (Figure 1B). Similar results were obtained when using AUC01 (Figure S1D), when considering only epitopes with at least 200 TCRs (Figure S1E) or when using as negatives in the test set TCRs found to interact with other epitopes ^26^ (Figure S1F) (see Materials and Methods). Predictions with TEMPO were also quite robust to contamination levels of up to 50% in the training data ^21^ (Figure S1G). Using a uniform baseline for renormalization in TEMPO led to significantly lower prediction accuracy (Figure S1H), confirming the importance of a robust model of baseline TCR repertoires to accurately capture TCR-epitope recognition specificity.

To benchmark TEMPO predictions with existing tools in their published version, we used the IMMREP23 ^14^ and the ePytope-TCR ^15^ studies (see Materials and Methods). All data related to these studies were excluded from the training of TEMPO. Here again, we observed similar or better AUC with TEMPO (Figure 1C&D, Figure S1I&J). Overall, these results demonstrate that TSPs accurately capture critical information underlying TCR-epitope recognition specificity available in today’s data.

In terms of running time and memory usage, TEMPO can score a million TCRs in one minute on a single standard CPU (Figure S1K), and requires approximately 100 times less disk space than tools like MixTCRpred ^12^.

### TSPs reveal key determinants of T cell epitope recognition specificity

To investigate how different features in TCR sequences contribute to epitope recognition specificity, we trained different versions of TEMPO, combining different features (i.e., V, J, CDR3 lengths and CDR3 sequences) or using each of them individually (Figure 2A and Figure S2A-D). The best performance was achieved when considering all features. The difference with a model considering only V and J genes was statistically significant, although modest in absolute numbers. When looking at single features, we observed that the most predictive one was V usage (Figure 2A and Figure S2A). This is consistent with the TSPs shown in Figure 1A and Figure S1C. To investigate the structural basis of these observations, we collected available X-ray structures of TCR-epitope complexes and annotated all residues in CDR loops as being encoded in the V segment (i.e., CDR1, CDR2 and beginning of CDR3), the J segment (i.e., end of CDR3) or resulting from insertions at the VJ junction in the CDR3 (referred to as ‘CDR3_VJ’) (see Materials and Methods). Counting the number of TCR residues contacting the epitope (i.e., peptide + MHC) revealed a much higher number of TCR residues encoded in the V and J segments than those in CDR3_VJ (Figure 2B, Table S3A). Even when restricting to contacts with the peptide, we still observed a significantly higher number of TCR residues encoded in the V and J segments than those in CDR3_VJ (Figure 2C, Table S3B). These results are fully consistent with our observation that V/J usage encodes much of the specificity in TCR-epitope recognition.

**Figure 2:**
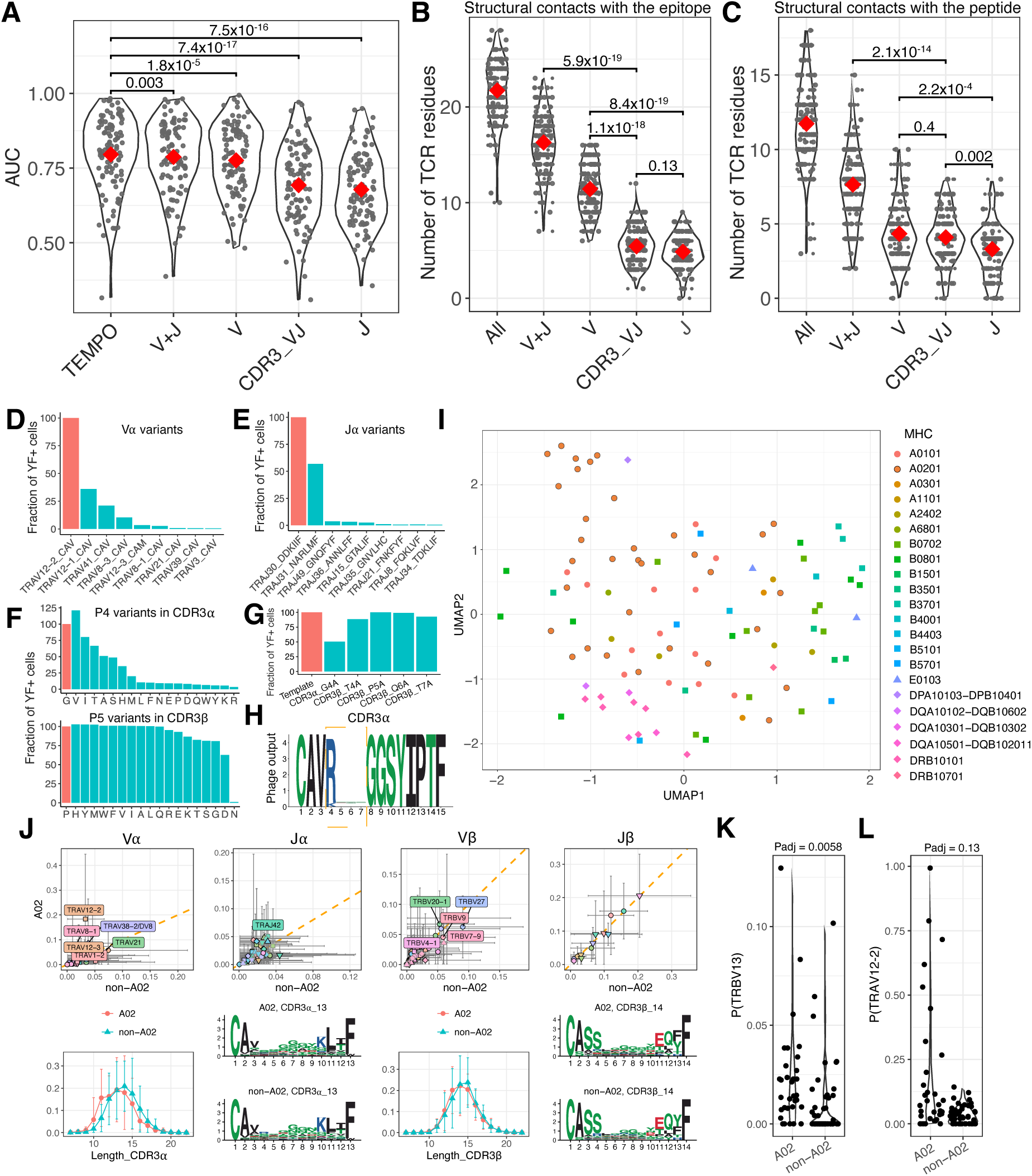
TSPs reveal key determinants of TCR-epitope recognition specificity. **A**: AUC for the cross-validation analysis with TEMPO considering different features for both chains. “CDR3_VJ” stands for the model based on CDR3 motifs (𝑃(𝐶𝐷𝑅3|𝐿)) with the 𝑄_P_(V,J|𝐿) normalization. Chain indication is omitted for clarity. **B**: Number of TCR residues in contact with the epitope (i.e., peptide + MHC) and encoded by the V and J segments or resulting from insertions at the V/J junction in CDR3s (referred to as “CDR3_VJ”). Data for X-ray structures with the same epitope have been averaged. P-values are indicated between statistically independent features only. **C**: Number of TCR residues in contact with the peptide and encoded by the V and J segments or resulting from insertions at the V/J junction in CDR3s. Data for X-ray structures with the same epitope have been averaged. P-values are indicated between statistically independent features only. **D**: Binding between the YF epitope and different Vα variants of TCR_YF. The y-axis shows the fraction of Jurkat cells transfected with different TCRs that were stained by the YF multimer. **E:** Binding between the YF epitope and different Jα variants of TCR_YF. **F:** Binding between the YF epitope and CDR3 variants of TCR_YF comprising all amino acids except cysteine at P4 in CDR3α (upper) and at P5 in CDR3β (lower). **G:** Binding between the YF epitope and CDR3 variants of TCR_YF with an alanine at CDR3 positions not determined by V/J usage and with no alanine in TCR_YF (P4 in CDR3α and P4-P7 in CDR3β, data for P4 in CDR3α and P5 in CDR3β are redundant (i.e., technical replicates) with those in panel F). **H:** CDR3α motif of TCRs obtained from a phage library with diversified CDR3α positions P4-P7 (orange dashed box) and screened against the NY-ESO-1 epitope (A0201_SLLMWITQC). **I:** UMAP view of epitopes with human TCRs embedded in their TSP space. Colors and shape indicate different MHC restrictions. **J:** Comparison of the average TSP of HLA-A*02:01 and non-HLA-A*02:01 restricted class I epitopes. **K:** TRBV13 usage in HLA-A*02:01 and non-HLA-A*02:01 restricted class I epitopes. **L:** TRAV12-2 usage in HLA-A*02:01 and non-HLA-A*02:01 restricted class I epitopes. P-values in panels A,B,C are computed with paired Wilcoxon test. P-values in panels J and K are computed with unpaired Wilcoxon test including multiple testing B&H correction.

To experimentally validate the importance of V and J usage in TSPs, we used a TCR binding to the YF epitope (TCR_YF: TRAV12-2, TRAJ30, CAVGDDKIIF, TRBV28, TRBJ2-7, CASTPQTAYEQYF) and replaced TRAV12-2 by different Vα genes with similar CDR3α sequences. Jurkat cells were transfected with the TCR variants and stained with the YF multimer (Figure 2D, Figure S2E, Table S4, see Materials and Methods). The results confirmed the high specificity for Vα usage observed in Figure 1A. We then replaced TRAJ30 by several Jα genes compatible with CDR3α of length 10. The results broadly confirmed the specificity observed in the TSP (Figure 2E, Table S4). Similar results were obtained with a TCR binding to A0201_GILGFVFTL (Figure S2F, Table S4). These results are consistent with the high sequence diversity of V (including in CDR1 and CDR2) and J gene sequences.

We then explored the limited specificity at some CDR3 positions when comparing to baseline CDR3 motifs with the same V/J usage. To this end, we selected the fourth position (P4) in the CDR3α and the fifth position (P5) in the CDR3β of a YF specific TCR and tested all amino acids except cysteine. In more than one third of the cases for P4 in CDR3α and in almost all cases for P5 in CDR3β the binding was partly or fully retained (Figure 2F, Table S4). We then replaced with alanine all CDR3 residues that were not encoded in V/J gene usage and were not themselves an alanine (i.e., P4 in CDR3α and P4-P7 in CDR3β). Here again, we observed that the binding was retained (Figure 2G). Overall, these few examples confirm the high specificity in V/J usage and limited specificity at CDR3 positions resulting from insertions at the V/J junction for TCRs recognizing the YF epitope.

To further investigate specificity (or lack thereof) in CDR3 positions not determined by V/J usage, we performed a phage display experiment with a TCR (alpha chain: TRAV21, CAVRPTSGGSYIPTF, TRAJ6) recognizing the NY-ESO-1 epitope (A0201_SLLMWITQC). Using our model of baseline CDR3α motifs (i.e., 𝑄(𝐶𝐷𝑅3_α_|𝑉_α_, 𝐽_α_, 𝐿_α_)) we observed that positions 4-7 displayed variability in baseline TCRα repertoires with the corresponding Vα and Jα (Figure S2G). These positions were diversified, and the resulting phage library was selected against the A0201_SLLMWITQC monomer (see Materials and Methods and Table S5). Unlike for the CDR3β which displayed clear specificity (see motifs in ref. ^27^), no specificity was observed at positions 5 to 7 (Figure 2H and Figure S2H), This is fully consistent with the limited interactions with the epitope mediated by the sidechains of these residues in X-ray structures of this TCR-epitope complex (Figure S2I). The arginine enriched at position 4 corresponds to the last amino acid of TRAV21. This unbiased phage display analysis provides a clear example of limited specificity encoded in some CDR3 residues not determined by the V/J choices.

We then explored whether some of the patterns observed in TSPs, including V/J usage, may reflect specificity for MHC alleles irrespective of the peptide. To this end, we considered all epitopes with at least 20 human TCRα and TCRβ and with documented MHC restriction. These epitopes were embedded in their TSP space and UMAP was used to visualize the data (see Materials and Methods). The results revealed some trends but also extensive variability, with no clear cluster encompassing all epitopes restricted to the same MHC (Figure 2I). To further investigate the impact of MHC restriction, we capitalized on the presence of many HLA-A*02:01 restricted epitopes (roughly one third of epitopes with human TCRs in our data) and compared the average TSP of HLA-A*02:01 versus non-HLA-A*02:01 restricted class I epitopes (Figure 2J) (see Materials and Methods). We observed some differences, but most of them were comprised within the expected variability across epitopes. Only one gene (TRBV13, enriched in HLA-A*02:01 restricted epitopes) reached statistical significance after multiple testing correction (Figure 2K). For TRAV12-2, which has been associated with several HLA-A*02:01 restricted epitopes ^25,28^, we could see over-representation in one third of the HLA-A*02:01 epitopes, but no enrichment in the remaining two thirds (Figure 2L). Overall, these data reveal substantial variability in V/J usage across epitopes restricted to the same MHC allele and only limited influence of shared MHC restriction on TSP similarity.

We next investigated the performance of TEMPO when trained on single-chain TCRs. Lower and roughly equal AUC values were obtained when considering only the alpha or only the beta chain (Figure S2J), which is consistent with previous studies ^29^. This indicates that, on average, both chains encode similar amount of specificity, which is also reflected in the number of TCR residues in each chain contacting the epitope or the peptide (Figure S2K and S2L).

### TSPs predict cross-reactivity

We hypothesized that similarity in TSPs may be used to predict cross-reactivity. To demonstrate this hypothesis, we experimentally quantified cross-reactivity and TSP similarity between the YF epitope and dozens of variants of this epitope (Figure 3A, Table 1).

**Figure 3:**
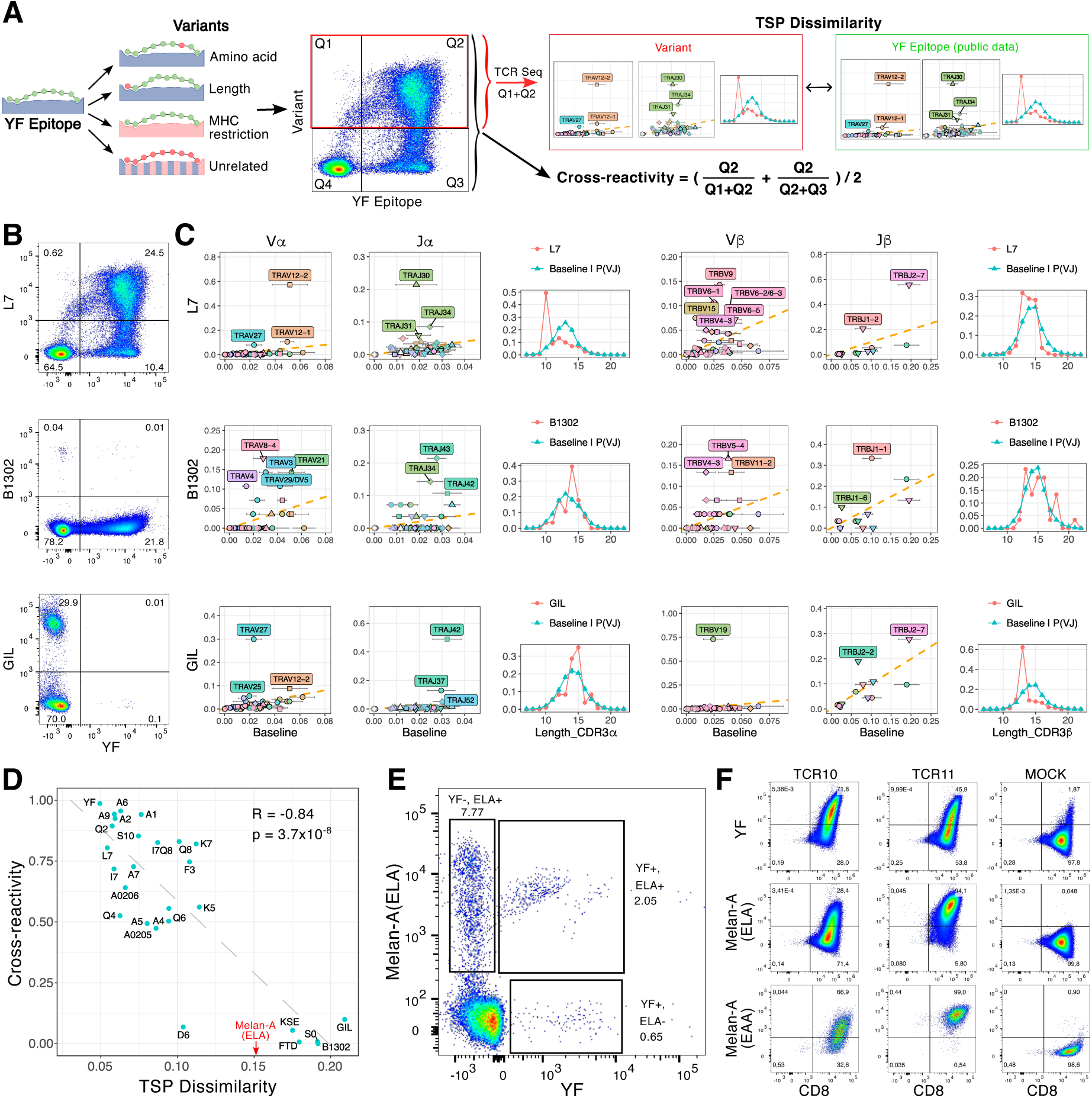
TSPs predict cross-reactivity. **A**: Overall pipeline to study the relationship between TSP dissimilarity and cross-reactivity. **B**: Example of cross-reactivity plots between the YF epitope (x-axis) and a sequence variant (L7), an MHC variant (B1302) or an unrelated epitope (GIL). **C**: TSPs for the three epitopes of panel B. **D**: Relationship between cross-reactivity and TSP dissimilarity. The red arrow shows the TSP dissimilarity between YF and Melan-A (ELAGIGILTV). **E:** Cross-reactivity between the YF and Melan-A (ELAGIGILTV) epitopes. **F:** Validation of the binding between the YF and Melan-A epitopes of two cross-reactive TCRs transfected in Jurkat cells and stained with the YF and Melan-A (i.e., ELAGIGILTV and EAAGIGILTV) multimers (see Table S9).

**Table 1:**
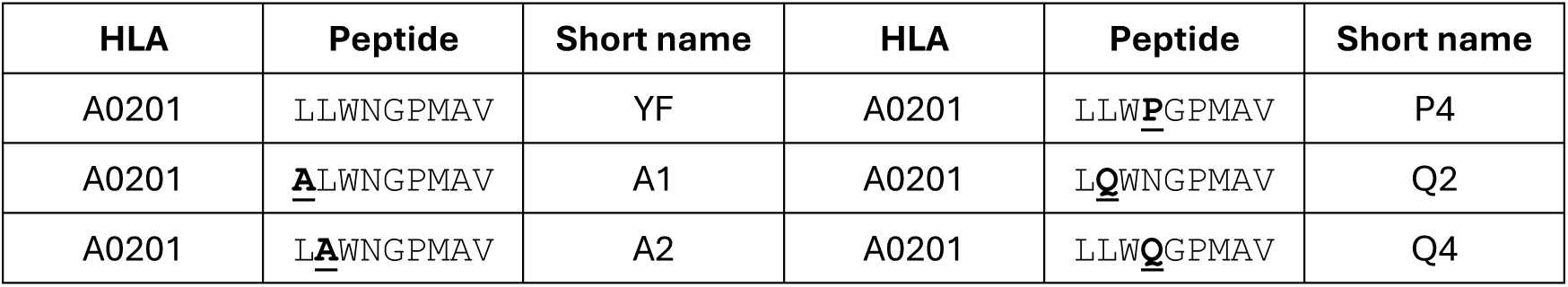
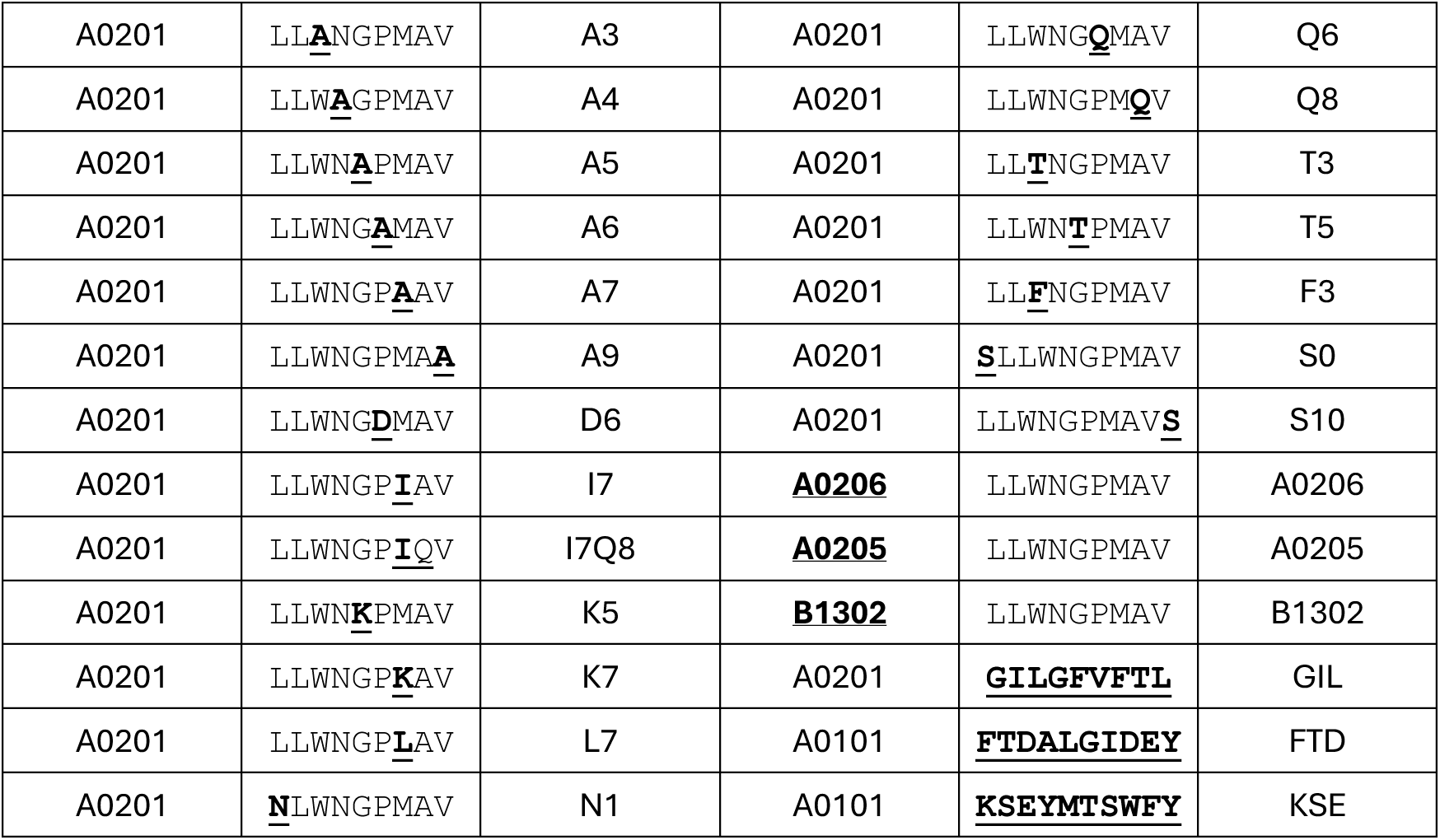
List of YF epitope variants and unrelated epitopes.

To start with, we stimulated Peripheral Blood Mononuclear Cell (PBMC) samples from three vaccinated donors with the YF peptide, sorted epitope-specific T cells with YF multimers and sequenced their TCRs with the SEQTR protocol ^30^ (see Materials and Methods, Table S6). The TSPs obtained in these samples displayed high similarity with the one derived from publicly available data (Figure S3A). For one sample, we sequenced the YF specific TCRs with the 10X Genomics single-cell TCR-seq pipeline (see Materials and Methods and Table S7). This revealed the presence of more than 400 distinct epitope-specific clonotypes. The TSP demonstrated high concordance with the one obtained with SEQTR (Figure S3B). We then compared the TSP obtained by sorting stimulated CD8 T cells based on the expression of the activation marker CD137 to the TSP obtained by multimer sorting. Here again, this resulted in a highly similar TSPs (Figure S3C). These results demonstrate that TSPs are reproducible across donors, TCR sequencing approaches and sorting strategies (i.e., binding / activation).

We next stimulated the same PBMC samples with different variants of the YF peptide, including all single alanine variants, other single or double variants, variants in peptide length, variants in MHC restriction, as well as some unrelated antigenic peptides (Table 1). CD8 T cells were stained with multimers corresponding to the YF epitope and all other epitopes to measure cross-reactivity (see Materials and Methods) (Figure 3B and Figure S4, Table S8). For each epitope for which we could detect some epitope-specific CD8 T cells, multimer+ cells were sorted and their TCRs were sequenced to determine the TSPs (see Figure 3C and Figure S5, Table S6, or the web interface https://tcrmotifatlas.unil.ch/Browse_epitopes_YF). Dozens of TCRs binding to diverse variants were successfully tested *in vitro* to confirm the results of the sequencing (Table S9, Figure S6, see Materials and Methods). We then developed an information theory-based measure of TSP dissimilarity (see Materials and Methods). To mimic a realistic situation where TCRs are collected across different donors with different sequencing protocols and in different places, publicly available data were used to derive the TSP for the YF epitope and the data generated in this work were used to establish the TSPs for the other epitopes. We observed a significant correlation between cross-reactivity and TSP dissimilarity (Figure 3D). Similar results were obtained with different variants of the TSP dissimilarity definition (Figure S7A). These results demonstrate that TSPs measured across different samples, protocols and places can be used to assess cross-reactivity between epitopes.

To further support this idea, we computed the TSP similarity between the YF epitope and all other epitopes with publicly available TCRs. The epitopes with the most similar TSPs were all related to the Melan-A antigen (A0201_ELAGIGILTV, see Figure 3D), which reflects among else a similar Vα usage (Figure S7B). FACS analysis of a PBMC sample confirmed some cross-reactivity between the YF and the Melan-A epitope (Figure 3E). TCRs from cross-reactive cells were sequenced and Jurkat cells were transfected with some of them and stained separately with the YF and Melan-A (i.e., both ELAGIGILTV and EAAGIGILTV) multimers to confirm the cross-reactivity (Figure 3F, Figure S7C and Table S9).

Overall, our results obtained with pools of thousands of native T cells, including hundreds recognizing the YF epitope, demonstrate that TSP similarity between epitopes can be used to predict cross-reactivity.

### TSPs reveal how TCR specificity evolves with epitope sequences, binding modes and MHC restrictions

Our set of TCRs recognizing multiple sequence, length and MHC restriction variants provides a unique opportunity to explore molecular mechanisms underlying TSP conservation. We first observed that amino acid variants at the beginning or the end of the YF peptide and preserving the binding to HLA-A*02:01 tend to preserve TSPs and lead to extensive cross-reactivity (Figure 4A). Conversely, several variants in the middle of the peptide significantly decreased TCR recognition. To functionally validate and extend these observations to more variants, we selected three TCRs binding to the YF epitope and tested recognition with all 162 single amino acid variants (i.e., 9×18, representing all but the original amino acids and cysteine at each position) using the X-scan assay (see Materials and Methods, Figure 4B, Figure S8A, Table S10). We observed that most variants at the N-terminus (P1&P2) or the C-terminus (P8&P9) and compatible with HLA-A*02:01 binding could be recognized, while many variants at middle positions (P3-P7, and especially P3-P5) impacted TCR recognition.

**Figure 4:**
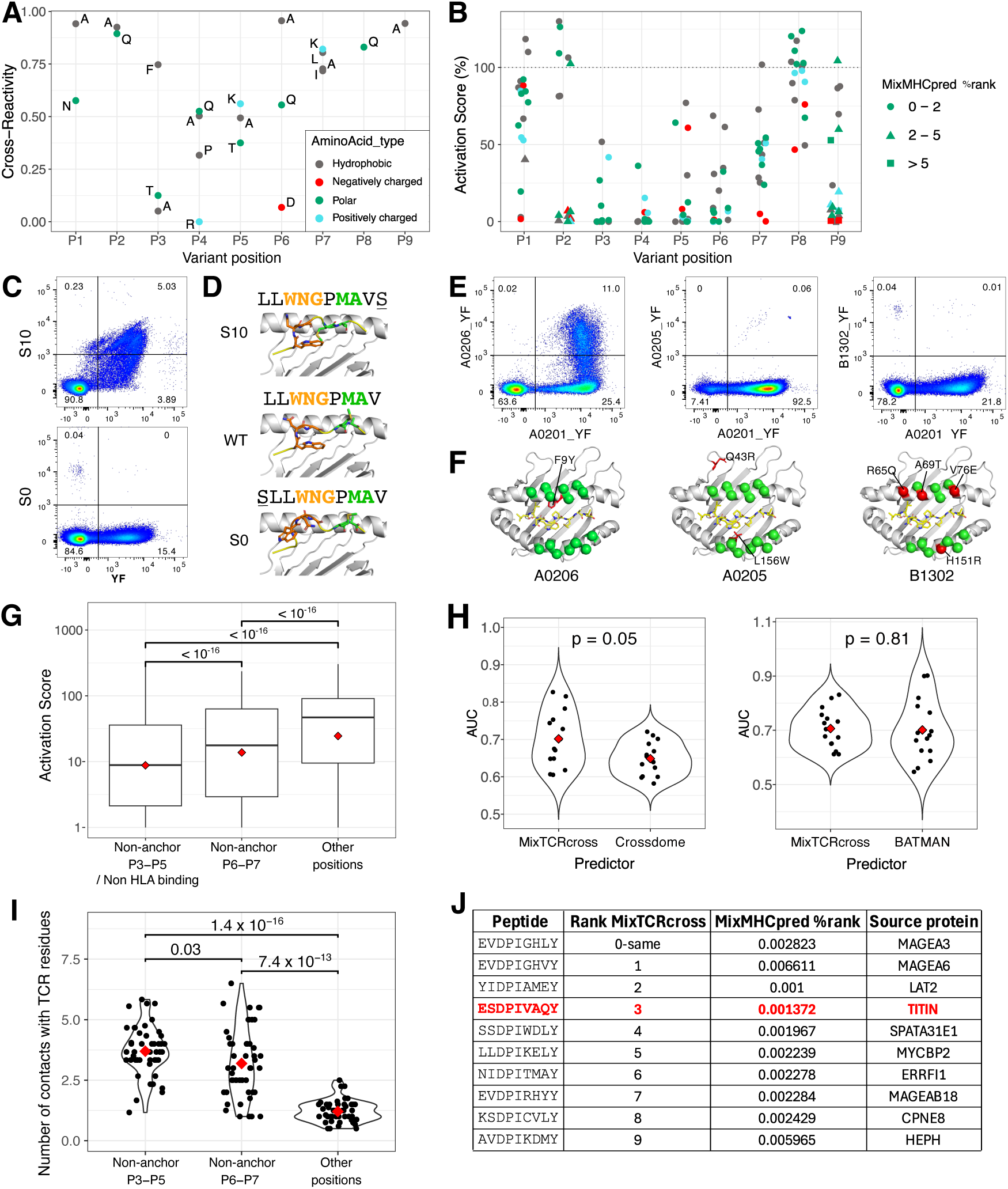
TSPs reveal how TCR specificity evolves with peptide sequences, binding modes and MHC restrictions. **A**: Summary of the cross-reactivity for the different YF sequence variants in Table 1. **B**: Results of X-scan assay for all single amino acid variants of the YF epitope. Each point represents the average reactivity of three different YF-specific TCRs. **C**: Cross-reactivity observed between the YF epitope and the two length variants (S0 and S10). **D**: X-ray crystallography of the S10 variant, the YF epitope and the S0 variant in complex with HLA-A*02:01. **E:** Cross-reactivity observed between the YF epitope and three MHC allele variants. **F:** Structural view of the conservation of the TCR interface for the three MHC alleles of panel E with respect to HLA-A*02:01. Residues in the TCR interface are shown with balls. Non-conserved residues are shown in red. **G**: Activity values for the single amino acid variants in the BATCAVE database grouped based on three different categories of positions used in MixTCRcross. **H**: Comparison of MixTCRcross with those of CrossDome and BATMAN in the unseen epitope context based on the 9-mer class I epitopes in the BATCAVE databases. **I**: Number of interactions with TCR residues mediated by peptide residues at the three different categories of positions used in MixTCRcross. **J**: Top 9-mer peptides from the human proteome most likely to show cross-reactivity with MAGE-A3 based on MixTCRcross predictions.

The two length variants (i.e., S0: SLLWNGPMAV and S10: LLWNGPMAVS) exhibited distinct behaviors. S10 showed high cross-reactivity and high TSP similarity with YF, while S0 displayed no cross-reactivity and had a distinct TSP (Figure 4C, see also Figure 3D and Figure S5). To determine the exact binding mode of these two peptides, we used X-ray crystallography (see Materials and Methods, Table S11). Both variants adopted a bulging binding mode. S10 had a structurally conserved N-terminal part with most structural rearrangements taking place after P6, while S0 displayed extensive structural rearrangements in the N-terminal part (Figure 4D, Figure S8B). The high cross-reactivity and high TSP similarity between YF and S10 is therefore consistent with our previous observations that changes outside of positions P3-P5 are more likely to preserve recognition of YF variants.

Regarding the impact of MHC restriction, we observed cross-reactivity and high TSP similarity for the YF peptide in complex with HLA-A*02:01, HLA-A*02:06 and to a lesser extent with HLA-A*02:05 but no cross-reactivity and a distinct TSP with HLA-B*13:02 (Figure 4E, see also Figure 3C-D and Figure S5). To interpret these results, we retrieved the MHC residues known to be involved in the TCR interface (see Materials and Methods). We observed that HLA-A*02:06 and HLA-A*02:05 had the same TCR interface as HLA-A*02:01, while HLA-B*13:02 displayed multiple amino acid changes (Figure 4F). For HLA-A*02:05, some differences were observed at positions that may indirectly impact TCR recognition (e.g., L156W, which may displace slightly the alpha helix). This likely explains the lower cross-reactivity.

Putting all these data together, our results support a cross-reactivity model for YF variants, called MixTCRcross, where presentation on MHC alleles with conserved TCR interface and conservation of non-anchor residues at positions P3 to P7 (e.g., variants at P1-P2 or P8-P9 and preserving MHC binding) in general leads to similar TSPs and high risk of cross-reactivity (see Materials and Methods). Among other variants of the YF epitope, those with conserved non-anchor residues at P3 to P5 show highest risk of cross-reactivity.

To explore whether MixTCRcross could show some predictive power for other epitopes, we collected data from the BATCAVE database^31^, which comprises thousands of variants of 15 class I epitopes recognized by more than 50 TCRs (see Materials and Methods). We observed a significantly lower activity for variants at non-anchor positions P3-P5 compared with those at non-anchor positions P6-P7, and even more with those at other positions (Figure 4G). We then compared MixTCRcross predictions with those from CrossDome ^32^ and from BATMAN ^31^ (see Materials and Methods). Similar or better predictions were obtained with MixTCRcross (Figure 4H). These results show that our proposed model captures in a fully interpretable way some important determinants of TCR specificity conservation propensity that are shared across multiple epitopes.

To investigate the structural basis of the MixTCRcross model, we computed for each peptide residue the number of contacts with the TCR across existing TCR-epitope structures (see Materials and Methods, Table S12). We observed that peptide residues at positions predicted by MixTCRcross to impact TSPs (i.e., non-anchor P3-P5, and to a lesser extent P6-P7) were involved in a significantly higher number of interactions with the TCRs compared to peptide residues at other positions (Figure 4I).

We next explored whether the MixTCRcross model could recapitulate cases of cross-reactivity with cancer epitopes. To this end, we considered the HLA-A*01:01 restricted MAGE-A3 epitope (`EVDPIGHLY`). We collected all 9-mers from the human proteome (1.1×10^7^ peptides in total). Ranking these peptides based on MixTCRcross predictions and excluding the MAGE-A3 peptide, we observed that the third hit corresponded to the TITIN peptide (`ESDPIVAQY`) responsible for the lethal cross-reactivity observed in the clinical trials with MAGE-A3-specific TCRs ^33^ (Figure 4J).

Altogether, our results reveal that some patterns are conserved across several epitopes to predict variants more likely to preserve TSPs, and therefore at higher risk of cross-reactivity, with key features including structural positions of the non-conserved residues and presentation on MHC alleles with similar TCR interfaces.

### TSPs enable interpretation of machine learning tools and reveal how AF3 can be used to decipher key determinants of TCR specificity

We hypothesized that TSPs may help interpreting machine learning TCR-epitope predictors. To this end, we scored a repertoire of 10^6^ TCRs with several tools for different epitopes (see Materials and Methods). TSPs built from the top 0.1% predicted TCRs were compared with the experimental TSPs. For epitopes with abundant training data, we observed that TEMPO and NetTCR could very accurately recapitulate the experimental TSPs (Figure 5A and S9A-B). MixTCRpred failed to recover Jα usage, while TULIP failed to recover Vα usage. The latter can be understood since TULIP only considers CDR3 sequences as input features, and several V genes have quite similar sequences in the CDR3 region (e.g., CAV in the Vα shown in Figure 5A for TULIP). Similar results were obtained with the GIL epitope (Figure S9B). We next took advantage of the new TSPs for the MHC variant (HLA-B*13:02) of the YF epitope and tested algorithms that in principle can make predictions for any epitope and any MHC restriction. In both cases, NetTCR and TULIP failed to reproduce the experimental TSP (Figure 5B and Figure S9C), although TULIP could in theory distinguish different MHCs. This confirms the low predictive power of machine learning approaches for ‘unseen’ epitopes ^14,15,34^, even when these epitopes differ only in their MHC restriction from epitopes with abundant training data.

**Figure 5:**
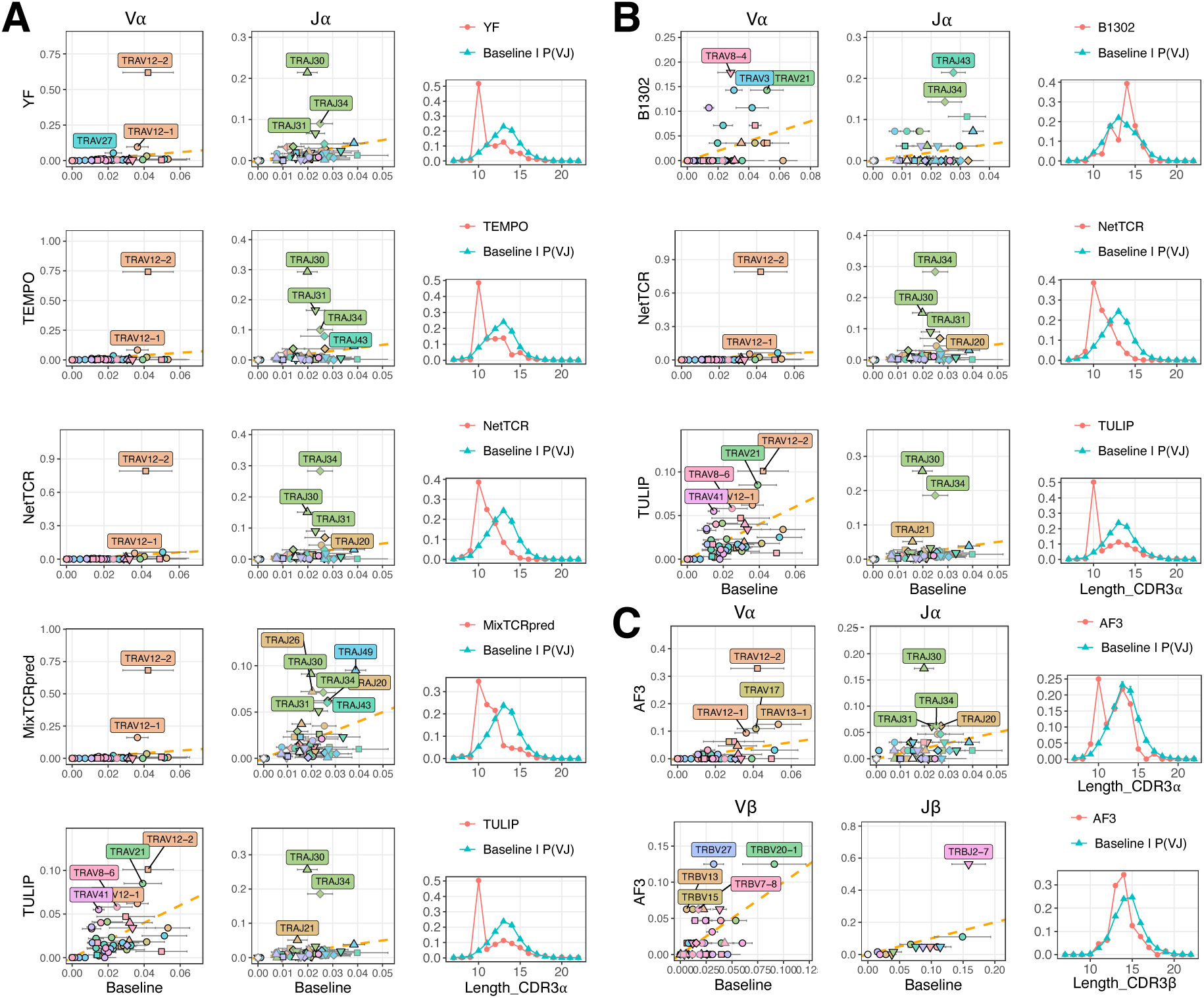
TSPs enable interpretation of machine learning tools and reveal how AF3 can be used to decipher key determinants of TCR specificity. A: Comparison between experimental and predicted TSPs for the YF epitope. **B:** Comparison between experimental and predicted TSPs for the B1302_LLWNGPMAV epitope. **C:** TSP predicted by AF3 for the YF epitope.

We then investigated whether AF3 ^16^ could be used to infer TSPs. As this tool cannot easily scale to a million TCRs, we sequenced with the 10X Genomics single-cell TCR-seq pipeline the *ex vivo* TCRαβ repertoire from a donor (Table S13) and ranked these TCRs based on their predicted binding to the YF epitope (i.e., interchain Predicted Alignment Error (PAE) score, see Materials and Methods). The TSP of the top predicted binders recapitulated several patterns found in the experimental TSP, including the enrichment for TRAV12-2, for several Jα genes and for CDR3α of length 10 (Figure 5C, Figure S9D). These results reveal how AF3 can be used to approximate TSPs even for epitopes without experimental TCR-epitope structural templates.

We next explored whether TSPs can reveal specific patterns in negative data used to benchmark TCR-epitope interaction predictors. Because of the lack of experimentally determined negatives, a widely used approach consists of using TCRs binding to other epitopes as negatives ^14,26^. We hypothesized that this may create specificity patterns reflecting TCRs binding to widely studied epitopes, such as GIL. To verify this hypothesis, we show the average TSP for TCRs used as negatives across all non-GIL epitopes from the IMMREP23 competition ^14^. A clear footprint of the GIL TSP could be observed (Figure S10A). These results indicate that using as negatives TCRs binding to other epitopes can create global biases which do not reflect realistic repertoires from patients.

We then hypothesized that TSPs may be used to detect low quality epitope-specific TCR datasets. We first selected one epitope from the 10X Genomics dataset previously suggested to contain many contaminants (A0301_KLGGALQAK) ^12,35^. The TSP showed very limited specificity (Figure S10B). We next explored whether some other epitopes widely used in different studies displayed such limited specificity. One of the most striking examples was A0201_NLVPMVATV (Figure S10C). This suggests that several TCRs reported to bind to A0201_NLVPMVATV may be false-positives.

Overall, these results show how TSPs can be used to assess the successes and limitations of machine learning TCR-epitope recognition predictions, explore TCR specificity based on AF3 predictions and investigate putative biases or data quality issues in epitope-specific TCRs.

## Discussion

Systematic understanding of the principles governing TCR-epitope recognition specificity has been hampered by the size and complexity of the TCR sequence space and the flexibility of the TCR-epitope interface. Here we show that, by using the natural alphabet of TCRs (i.e., V and J genes, CDR3 lengths and sequences) and carefully modelling baseline TCR repertoires, we can address this complexity in a fully interpretable way which accurately captures key determinants of TCR-epitope recognition specificity encoded in today’s data across hundreds of epitopes. The conservation of TSPs across donors, TCR-sequencing protocols and sorting strategies demonstrate that this framework is robust to various types of biases or batch effects.

Our results indicate that much of the specificity in epitope recognition by native TCRs is encoded in V/J usage, with many epitopes showing up to ten-fold enrichment in specific V or J genes. This is consistent with previous observations on a few selected epitopes ^5,25,36–38^ and structural data where extensive contacts with epitope residues are observed in the parts of the TCR sequences determined by the choices of V and J genes (i.e., CDR1, CDR2, beginning and end of CDR3). Beyond specificity in V/J usage, TSPs can be used to quantify additional specificity in CDR3 lengths (e.g., the preference for CDR3α of length 10 in YF-specific TCRs) or CDR3 residues that are not determined by V/J gene choices (e.g., the known “RS” motif in CDR3β of GIL-specific TCRs, see Figure S1C and other examples at https://tcrmotifatlas.unil.ch). Our work further reveals high variability in V/J usage across epitopes, including those restricted to the same MHC allele. Apart from a few exceptions (e.g., YF and Melan-A), this indicates that the rules of TCR-epitope recognition are distinct for most epitopes.

The relatively limited specificity observed in some (though not all) CDR3 binding motifs when regressing out the influence of V/J usage (i.e., by comparing to CDR3 motifs from baseline repertoires with the same V/J usage) does not mean that CDR3 sequences do not contribute to TCR-epitope recognition. Instead, it reflects that a substantial fraction of the CDR3 residues (including many that directly interact with the epitope) is encoded in the choice of V and J genes.

TSPs consider each feature (i.e. V/J genes, CDR length and CDR3 amino acids) as independent and therefore cannot capture correlations. Such correlations exist, but many are unrelated to epitope recognition. For instance, extensive correlations exist between Vα and Jα, as a result of constraints in V/J recombination ^39^. Correlations related to epitope recognition specificity can arise from distinct clusters of TCRs recognizing the same epitope. Evidences of such clusters have been reported ^5,6,40^. However, it remains unclear how much machine learning tools benefit from modelling such correlations. Moreover, the presence of contaminants can result in apparent clusters that do not reflect epitope recognition specificity ^21^. Our demonstration that a linear model based on TSPs reaches similar or better accuracy than other state-of-the-art machine learning tools indicates that neglecting such correlations is a reasonable compromise between accuracy and interpretability. The limited impact of disregarding correlations can be further understood by observing that non-binding combinations of features (e.g., V/J genes) compatible with TSPs make up only for a very tiny fraction of baseline TCR repertoires from patients (e.g., 𝑄(TRAV12-2, TRAJ34, 𝐿_α_=10, TRBV28, TRBJ2-7, 𝐿_*_=13) = 3 × 10^+,^ for the non-binding example in Figure 2E) and are rarely expanded. Our analysis of the recognition of multiple variants of the YF epitope did not rely on a list of pre-defined TCRs. This is the reason why we could assess not only the conservation but also the emergence of new TSPs in cases like B1302_LLWNGPMAV. Our results suggest a model for TCR specificity conservation propensity across YF variants which can be summarized with a few criteria. These include conservation of peptide residues at non-anchor middle positions, and conserved presentation on MHC alleles with the same TCR interface. The observation that this relatively simple model has some predictive power for other epitopes indicates that our proposed criteria capture, in a fully interpretable way, important determinants of TCR specificity conservation which are shared across many 9-mer class I epitopes and are consistent with structural contacts between peptide residues and TCRs, even if several exceptions exist ^1^. Epitopes that fulfill these criteria show high risk of cross-reactivity, as illustrated in the MAGE-A3 – TITIN example. For epitopes that do not pass these criteria, cross-reactivity is less likely, but cannot be excluded. In these cases, our work shows that reliable predictions of cross-reactivity should leverage TSP similarity. An alternative is to generate data for a TCR recognizing a given epitope and train tools which can learn cross-reactivity patterns related to a specific TCR-epitope pair beyond the shared patterns described in this work^31^.

Our approach to interpret machine learning TCR-epitope interaction predictors will help address some pitfalls of specific tools, such as limitations in predicting V usage when considering only CDR3 sequences. As TSPs do not require negative data (i.e., they use a statistical model of baseline TCR repertoires instead), this approach provides a fully interpretable alternative to measures like AUC that typically rely on artificially generated negatives which can display significant biases.

In this study, we mainly focused on epitopes with experimentally available TCRs. However, our investigation of TSP conservation and our observation that AF3 can, at least to some extent, approximate TSPs by scoring TCRs from *ex vivo* repertoires indicate that this approach may be extended to infer key determinants of TCR recognition for a much larger panel of epitopes.

Altogether, our work demonstrates that key determinants of TCR-epitope recognition specificity can be accurately captured and visualized with TSPs. This framework, together with the accompanying TCR Motif Atlas (https://tcrmotifatlas.unil.ch) and the TEMPO tool, enables investigation and interpretation of TCR-epitope recognition specificity, as well as efficient annotation of large TCR repertoires for many epitopes across different diseases. Our observation that AF3 can be used to approximate TSPs even in the absence of an experimental structural template provides a way to expand the investigation of TCR recognition specificity to many more epitopes. Finally, our proposed criteria to predict TCR specificity conservation could guide the ob-target toxicity assessment of TCRs considered for clinical applications.

## Materials and Methods

### Collection of baseline TCR repertoires

To build an accurate model of baseline TCR repertoires accounting for the variability across studies and individuals, data available at iReceptor were collected ^41^ (Table S1). These include paired and single-chain TCR repertoires from a large diversity of studies including many different sequencing protocols, many TCR reconstruction approaches, as well as different immunological status and genetic background of the donors. Samples from particular immunological conditions (e.g., leukemia) and showing large discrepancies in V/J usage were not included. Entries with V/J names that did not follow the IMGT reference were corrected based on an extensive manual compilation of all encountered alternative names. Cases where correction was not possible (e.g., TRBJ2S13) were not considered. Only TCRs with CDR3 of length 7 to 22 and with CDR3 sequences compatible with V and J gene annotation based on the first one and the last two amino acids were considered.

In total this led to a collection of more than 6 million TCRα and more than 200 million TCRβ sequences in human and mouse. For each study and for each chain, V/J usage, CDR3 length distributions and CDR3 amino acid frequencies for each length were computed. The means (i.e., 𝑄(𝑉), 𝑄(𝐽), 𝑄(𝐿) and 𝑄(𝐶𝐷𝑅3|𝐿), where the latter represents the 20 x *L* amino acid frequency matrix at each position of CDR3s of length *L*) and the standard deviations of these probability distributions were computed across all studies for each chain in each species (for clarity, chain and species information are omitted from the formula). Residual probabilities for the CDR3 length and amino acid usage for each VJ combinations (i.e. 𝑄(𝐿|𝑉, 𝐽) and 𝑄(𝐶𝐷𝑅3|𝑉, 𝐽, 𝐿)) were further derived.

### TSP construction

TSPs provide a fully interpretable framework to compare a set of epitope-specific TCRs (probabilities designed with the letter *P*) with a baseline TCR repertoire (probabilities designed with the letter *Q*) for both alpha and beta chains (for clarity, the chain is not included in the rest of this section). For V, resp. J, genes the y-axis shows 𝑃(𝑉), resp. 𝑃(𝐽), and the x-axis shows the mean and standard deviation of 𝑄(𝑉), resp. 𝑄(𝐽) (Figure 1A). For CDR3 length distributions, 𝑃(𝐿) is compared with the expected baseline distribution with the same V/J usage. This is mathematically defined as:

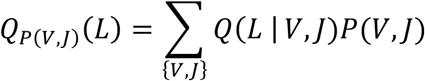

For CDR3 sequences of a specific length L, 𝑃(𝐶𝐷𝑅3|𝐿) represents the amino acid frequency at each position in the CDR3 of epitope-specific TCRs (i.e., 20 *x L* matrices), which are visualized as sequence motifs. The baseline motifs used for comparison are mathematically defined as:

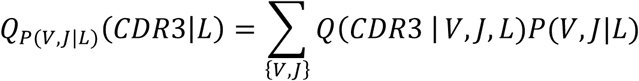

By default the CDR3 motifs corresponding to the most frequent CDR3 length (i.e., argmax{𝑃(𝐿)})) are shown in the main plot (Figure 1A), and motifs for other CDR3 lengths are shown in separate plots (examples in Figure S1B, see also https://tcrmotifatlas.unil.ch).

For visualization, TSPs display the names of specific V and J genes, based on their frequency and fold-change with respect to baseline repertoires. The web interface (https://tcrmotifatlas.unil.ch) further allows users to mouse over V/J genes to retrieve names, log fold-change, Z-scores and CDR3 sequences. By default, TSPs are shown at the gene level. Allele-level visualization is supported, although some care is needed when interpreting the results given that allele-level baselines are more challenging to accurately estimate.

### TCR-epitope motif-based interaction predictor (TEMPO)

TEMPO builds upon the widely used concept of binding motifs, which includes estimation of frequencies in input data (e.g., epitope-specific TCRs) and renormalization by baseline frequencies. The different coefficients of the predictor are computed as: 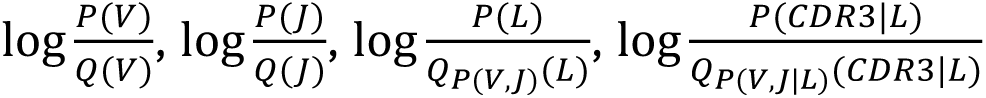 for both chains. These values are directly derived from the analytical solution of the log-likelihood maximization of a probabilistic model defined as 𝑃(𝑉, 𝐽, 𝐿, 𝐶𝐷𝑅3) = 𝑃(𝑉)𝑃(𝐽)𝑃(𝐿|𝑉𝐽)𝑃(𝐶𝐷𝑅3|𝑉𝐽𝐿). Regularization was done by adding a pseudocount of 50 in the epitope specific TCRs. The same pseudocount correction was performed on the baseline to ensure that features not observed in the epitope-specific TCRs and the baseline (e.g., non-cysteine amino acids at the first CDR3 position) will not contribute to the predictor.

The final score of a TCR is given by:

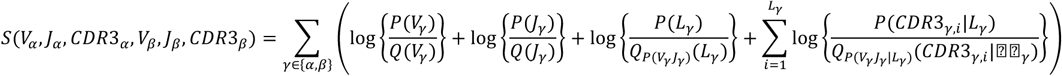

where 𝐶𝐷𝑅3_α,i_stands for the amino acid at the i^th^ position in CDR3α. To improve interpretability, %rank were computed by comparing the score of a TCR given in input with a distribution of scores coming from a dataset of 10^6^ TCRs sampled from baseline TCR repertoires.

### Collection of epitope-specific TCRs

TCR-epitope pairs were collected from different databases, including VDJdb and IEDB, as well as specific studies ^12,42,43^, in both human and mouse. All V and J names were mapped to the IMGT reference and all CDR3 were checked for compatibility with V/J genes based on the first one and last two amino acids. For each epitope, the MHC restriction was manually verified in the original publications. For clarity, epitopes are referred to by the MHC short name (e.g., A0201) and the peptide sequence (e.g., LLWNGMPAV), resulting in names like A0201_LLWNGPMAV. Cases where the MHC allele could not be retrieved (either because of the nature of the experiments or inability to retrieve it from the source publications) are denoted as ‘na’. Epitope peptide source organisms and source proteins were manually verified and harmonized across entries. Entries coming from the 10X Genomics dataset were not included, as this dataset was reported to contain a substantial amount of contaminants ^12,35^. Entries with incomplete chains (e.g., missing J segment) were not considered. Only TCRs from human or mouse were considered. This led to a total of 93,155 TCR-epitope pairs (Table S2). Among epitopes with at least 20 paired human αβTCRs, the most frequent allele was HLA-A*02:01 (33%, without counting the YF variants specifically designed in this work), which reflects more or less the frequency in the human population.

### Benchmarking predictions of TCR-epitope interactions

For the cross-validation analysis, all epitopes with at least 20 complete paired TCRαβ were considered. These TCRs were split into 5 subsets using iteratively one subset as test data and the other four as training data. To avoid leakage between training and testing data, we excluded from the training data any entry that had the same CDR3α or CDR3β as a TCR in the test set. For the negatives used in the test set, we selected TCRs from TCR repertoire studies not included in our baseline (see Table S1) and used 4 times more negatives than positives. For MixTCRpred ^12^, NetTCR ^11^ and TULIP ^8^, negatives in the training data were retrieved from the baseline TCR repertoires coming from the same studies as those used to estimate the baseline in TEMPO. TEMPO, MixTCRpred, NetTCR and TULIP were trained on the exact same data to ensure that differences in AUC could not result from differences in the training data of the different tools. NetTCR was downloaded from https://github.com/mnielLab/NetTCR-2.2 and trained using the train_nettcr_2_2_peptide.py script; TULIP was downloaded from https://github.com/barthelemymp/TULIP-TCR and trained with the full_learning_new.py script and the shallow0_decoupled.config.json configuration; MixTCRpred was retrained with our internal training module. AUC and AUC01 (i.e., partial AUC based on a threshold of 0.1 on the false-positive rate) metrics were used to assess the predictive power. Alternatively, negative data in the test set were generated by randomly selecting TCRs binding to other epitopes (the so-called “swapped negatives”), ensuring similar numbers of TCRs from each epitope (Figure S1F). Contaminants in the training data (Figure S1G) were simulated by adding randomly selected TCRs from the TCR repertoires considered in this work at different fractions of the number of positives in the training set.

To compare with existing publicly available methods in their published form, we capitalized on two recent benchmarking studies (i.e., IMMREP23 ^14^ and ePytope-TCR ^15^) whose data have not been used for training public TCR-epitope interaction predictors. All data from IMMREP23 itself, from the Immunoscape study ^44^ and from our own internal investigation of IMMREP23 epitopes ^14^, as well as any TCR present in these two benchmarking datasets set, were removed from the training of TEMPO and results were compared to those obtained with the published version of MixTCRpred (v1.0), NetTCR (v2.2) and TULIP (v1.0). Of note, in the IMMREP23 competition, MixTCRpred had been retrained with extended data, which is why AUC values are not the same here and in the IMMREP23 publication ^14^. For epitopes not supported in MixTCRpred and TEMPO, AUC and AUC01 were set to 0.5. For the uniform baseline, all values for the baseline distributions (𝑄) were set to the inverse of the alphabet size.

For the time benchmarking, TEMPO, MixTCRpred, NetTCR, and TULIP were tested on a standard workstation equipped with an Intel Core i9-11900H CPU (8 cores, 16 threads) and an NVIDIA GeForce RTX 3050 Laptop GPU (4 GB VRAM), running Linux with CUDA 12.2.

### Analysis of contacts with epitopes mediated by TCR residues

Experimental structures of human TCR-epitope complexes were retrieved from the Protein Data Bank (PDB) ^45^ [downloaded on 19.04.2024 for class I and on 25.08.2025 for class II]. Only X-ray structures with a resolution of 4 Å or better and class I MHC molecules were included. Redundant entries, defined as those with the same six CDR and epitope sequences, were filtered by retaining the structure with the highest number of contacts. Only TCRs with native V genes were considered (e.g., excluding cases with mutations in CDR1/2 enhancing the affinity). Contacts between a TCR residue and either epitope (i.e., peptide + MHC) or peptide residues were defined based on a threshold of 5Å between any heavy atom. TCR residues were annotated as encoded by the V segment if they belonged to CDR1, CDR2 or the part in the beginning of the CDR3 that matches the sequence of the corresponding V segment. TCR residues were annotated as encoded by the J segment if they belonged to the part in the end of the CDR3 that matches the sequence of the corresponding J segment. The other residues in the CDR3 were annotated as “CDR3_VJ”. The number of TCR residues in each category making contacts with the epitope (i.e., peptide+MHC) or the peptide only was computed for each structure (Table S3). To avoid biases due to well-studied epitopes with several X-ray structures, these numbers were averaged across structures corresponding to the same epitope.

### TCR Transfection into Jurkat Cells and peptide-MHC Multimer Staining

Paired TCR α and β chain sequences were obtained from GeneArt (Thermo Fisher Scientific) as gene strings. The DNA sequences were codon-optimized, and the human constant regions were replaced with mouse constant regions. These modified DNA fragments served as templates for in vitro transcription (IVT) and polyadenylation using the HiScribe™ T7 In Vitro Transcription Kit (New England Biolabs), following the manufacturer’s instructions. TCRα/β mRNAs were co-transfected into Jurkat cells using the Neon™ Electroporation System (Thermo Fisher Scientific) as described (PMID: 39276771). The Jurkat cells used were derived from the T Cell Activation Bioassay NFAT line (Promega) and had been genetically modified in-house via CRISPR/Cas9-mediated knockout of endogenous TCRα and TCRβ chains and by stable transduction with CD8a and CD8b (Jurkat-NFAT-CD8).

For electroporation, Jurkat cells were washed in PBS, resuspended at 20 × 10E6 cells/mL in Buffer R (Neon Transfection System 10 μL Kit), and 2 × 10E5 cells were co-electroporated with 300 ng of each TCR chain mRNA using the following parameters: 1325 V, 10 ms, 3 pulses. After electroporation, cells were resuspended in complete medium and incubated overnight at 37°C.

Each batch of TCR-electroporated cells was split in two. One half was stained with in-house-synthesized peptide-MHC multimers conjugated to PE, followed by a surface staining panel consisting of anti-human CD3 APC-Fire 750 (clone SK7, BioLegend), anti-human CD8 FITC (clone SK1, BioLegend), and Aqua viability dye (Thermo Fisher Scientific). The other half was stained with the same surface panel, including anti-mouse β chain APC (clone H57-597, BioLegend) to assess electroporation efficiency.

The template TCR binding to YF (TCR_YF) consists of TRAV12-2, TRAJ30, CAVGDDKIIF, TRBV28, TRBJ2-7, CASTPQTAYEQYF. The template TCR binding to A0201_GILGFVFTL (TCR_GIL) consists of TRAV27, TRAJ42, CAGGGSQGNLIF, TRBV19, TRBJ2-7, CASSIRSSYEQYF.

### Phage display

Phage libraries with diversified positions at P4-P7 in the CDR3α were constructed on the codon-optimized (*E. coli*) 1G4 single-chain TCR scaffold incorporating Vβ CDR2 “anchoring” point mutations (VGAGI > VSVGM; c50). These mutations have been shown to enhance TCR affinity without appreciably modifying the cognate TCR-peptide contacts or TCR docking orientation ^46^. The P4-P7 residues (‘RPTS’) were randomized in a 2-step PCR using primers spanning the signal peptide/Vα junction (NcoI/FOR) and the CDR3α (KpnI/REV), the latter containing trimer-defined (TRIM) codon mixtures (Ella Biotech GmbH, Germany) formulated to approximate the native CDR3α amino acid frequencies found at these positions. The diversified Vα fragment PCR product was double-digested with NcoI/KpnI and ligated into a pUC19-based phagemid (pCHV101) vector backbone housing the 1G4 Vβ[c50]Cβ to generate a library of TCR variants fused in-frame with the N-terminus of the M13/f1 gIII major coat protein. Phage libraries were produced from electroporated TG1 *E.coli* according to established protocols using monovalent M13 helper phage (Life Technologies). Phage panning was performed according to standard protocols against monomeric, biotinylated HLA-A*02:01_NY-ESO-1157-165 immobilized on M-280 streptavidin magnetic beads (Thermo Fisher Scientific) using an irrelevant HLA-A*0201 antigen for negative deselection of any non-selective binders. A single round of panning was conducted in which unbound phages were eluted following 5x PBST (PBS containing 0.1% Tween-20) and 1x PBS washes using trypsin cleavage and amplified by infecting logarithmic-phase *E. coli.* Following bulk purification of amplified output phagemid DNA, enrichment of specific CDR3α was assessed by NGS (2×300 bp paired-end) using an in-house Illumina MiSeq platform. Raw sequencing data were processed with MiXCR v3.0.13 with standard parameters (mixcr analyze amplicon -- species hs starting-material dna-- 5-end v-primers --3-end j-primers --receptor-type TRA) to reconstruct CDR3α sequences both for the input library and the output library after selection with NY-ESO-1 (Table S5). For the output library, MoDec ^47^ was used to remove putative contaminants. CDR3α motifs for the input and output libraries were built with ggseqlogo ^48^ (Figure 2H, see Figure S2H for motifs built without running MoDec).

### Embedding epitopes into TSP space

To embed epitopes into their TSP space, the TEMPO coefficients log 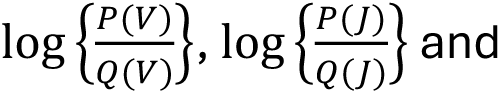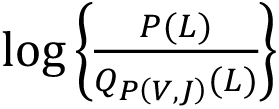 (including pseudocount regularization) for both chains were used as features. This resulted in a 220 dimensions space. UMAP (based on R package umap with default parameters) was used to visualize the data in 2D. All epitopes with a known MHC restriction and at least 20 human TCRα and TCRβ were used for this analysis.

### Average TSPs

To explore TSPs representing groups of epitopes (HLA-A*02:01 versus non-HLA-A*02:01 restricted class I epitopes), the average and standard deviation for each feature in the TSPs were computed across the epitopes in each group. For the non-HLA-A*02:01 restricted epitopes, an average TSP was first computed for epitopes restricted to each MHC. These TSPs were then averaged across all MHC alleles to avoid biases resulting from common MHC alleles with multiple epitopes. Only epitopes restricted to human class I alleles were considered.

### Collection of PBMC samples

Volunteers for blood donation were recruited from persons who had received YF vaccination due to future YF virus exposure risk and were HLA-A*02:01 positive ^25^. The study protocol was approved by the Human Research Ethics Committee of the Canton de Vaud (protocol 329/12). Upon written informed consent, eligible volunteers donated blood according to the standards of the Blood Transfusion Center (Service Vaudois de Transfusion Sanguine).

Peripheral blood samples were obtained in heparin-coated tubes and immediately processed for cryopreservation for downstream applications. To isolate PBMCs, heparinized whole blood was first diluted 1:1 with phosphate-buffered saline (PBS) and carefully layered onto Lymphoprep™ for density gradient separation. Samples were centrifuged for 30 minutes at 400 g at room temperature without break. The PBMC layer was collected and washed once in complete medium (RPMI 1640 supplemented with penicillin, streptomycin, HEPES buffer, and 10% fetal calf serum complete medium by centrifugation at 350 g for 5 minutes. Cells were then immediately resuspended in ice-cold freezing medium (CM with 40% FCS and 10% dimethyl sulfoxide), frozen overnight at −80 °C, and transferred into liquid N2 for long-term storage. In total, three donors were used (LAU5050, LAU5052, LAU5013).

### Stimulation and sorting of epitope-specific CD8 T cells

CD8+ T-cells from cryopreserved PBMC were enriched using MACS (Magnetic Activated Cell Sorting) technology (Miltenyi Biotec). The CD8 negative fraction of cells was irradiated (30 Gray) and used as antigen-presenting cells to stimulate peptide-specific T-cells. The CD8+ and irradiated CD8-cells were co-cultured together with the different peptides in RPMI supplemented with 8% human serum (Biowest), Penicillin/Streptomycin and beta-mercaptoethanol. Recombinant human IL-2 was added to the culture at a starting concentration at 10U/ml and increased gradually to 100U/ml by changing the culture medium every second day. After 11-14 days of culturing, T cells were analyzed by FACS with pMHC multimers to determine the cross-reactivity (YF multimer in APC and other multimers in PE). Epitope-specific CD8 T cells were then sorted and sequenced by SEQTR method for each epitope listed in Table 1 for which such cells could be detected. The same approach was used to explore cross-reactivity between YF and Melan-A (A0201_ELAGIGILTV).

### TCR sequencing and reconstruction

TCR sequencing of sorted CD8 T cells was performed with SEQTR pipeline for both alpha and beta chains ^30^. Raw sequencing data were processed with MiXCR to reconstruct both the alpha and beta chains. To account for expected batch effects, a dedicated baseline was built for SEQTR data, based on TCR repertoires from different PBMC samples ^30^.

### Single-cell TCR sequencing

Single-cell TCR sequencing was performed using the Protocol chromium GEM-X Universal 5’Gene Expression v3 4-plex OCM (CG000770 rev A) (10x Genomics) according to the manufacturer’s protocol. Pre-stimulated and tetramer-sorted T cells were washed, counted, and adjusted to the recommended loading density prior to microfluidic partitioning. Single cells were encapsulated with barcoded gel beads in GEMs (Gel Beads-in-Emulsion) using the Chromium X controller. Within each GEM, cell lysis and reverse transcription produced barcoded full-length cDNA, which was subsequently amplified. Amplified cDNA underwent targeted enrichment of T-cell receptor (TCR) α and β chain sequences via 2 PCR amplifications according to the 10x genomics workflow. A 5’ Gene Expression library was also made from the same cDNA. Libraries were indexed, pooled, and sequenced on the AVITI platform (Element Biosciences) on a Cloudbreak Freestyle flow cell with sequencing parameters 10-10-28-90 (index 1, index 2, read 1, read 2). Demultiplexing was done using the Bases2Fastq software (v2.0, Element Biosciences). Raw sequencing data were processed with Cell Ranger (v9.0.0, 10x Genomics) using the refdata-cellranger-vdj-GRCh38-alts-ensembl-7.1.0 genome reference, to assemble contigs, identify clonotypes, and annotate paired TCR α/β sequences, yielding high-resolution TCR repertoire profiles for downstream analysis.

### Cross-reactivity analysis

To measure cross-reactivity, CD8 T cells were stained with both the A0201_LLWNGPMAV multimer and the multimer corresponding to each variant. Cross-reactivity was assessed by the following formula ½ Q2/(Q1+Q2) + ½ Q2/(Q2+Q3), where Q2 denotes the fraction of double-positive cells, Q1 the fraction of cells positive only for the first epitope and Q3 the fraction of cells positive only for the second epitope (Figure 3A). In this way, fully cross-reactive epitopes have a score of 1, while non-cross-reactive epitope have a score of 0.

### Quantifying TSP dissimilarity

TSP dissimilarity for two epitopes (E1 and E2) was computed as:

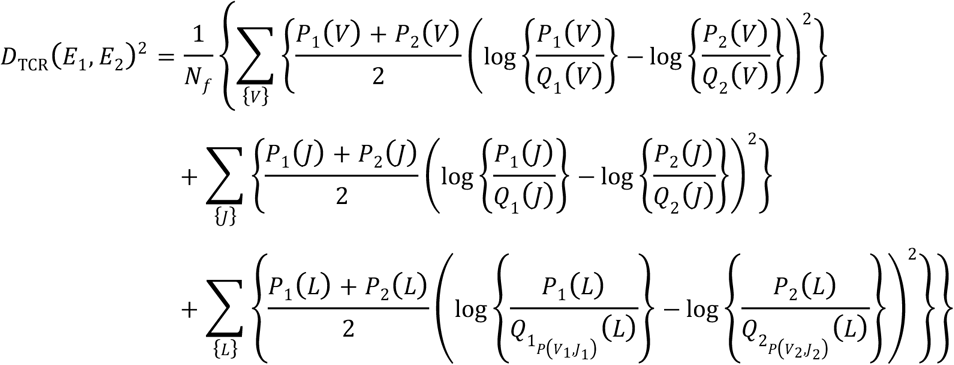

Where *Nf* stands for the number of features (i.e., number of V genes, J genes and number of CDR3 lengths). As before, for clarity, the sum over the two chains is omitted from the formula. The version including the CDR3_VJ contribution used in Figure S7A was obtained by adding the following contribution:

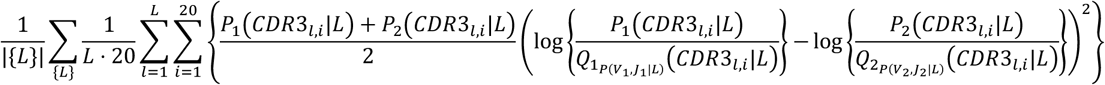

### X-scan assay

For each TCR (TCR_YF: TRAV12-2, CAVGDDKIIF, TRAJ30, TRBV28, CASTPQTAYEQYF, TRBJ2-7, TCR_YF2: TRAV12-2, CAVNPDKIIF, TRAJ30, TRBV4-2, CASSQEDRGPEKLFF, TRBJ1-4; TCR_YF3: TRAV12-2, CAAGDDKIIF, TRAJ30, TRBV29-1, CSVATSGGSNEQFF, TRBJ2-1), a dose–response experiment using the T Cell Activation Bioassay NFAT (Promega) was performed to determine the EC90. Briefly, 2×10E4 TCR-transfected Jurkat-NFAT-CD8 cells (as described above) in 25 μL were co-cultured with 4×10E4 T2 cells (ATCC) in 25 μL in a white 96-well plate (Costar) in the presence of a serial dilution of the YF native peptide (LLWNGPMAV) (25 μL). All conditions were tested in duplicate. After overnight incubation at 37 °C, the plate was equilibrated to room temperature, and one volume of Bio-Glo™ Reagent (Promega) was added to each well. Luminescence was measured using a Spark microplate reader (Tecan). Data were analyzed by nonlinear regression (four-parameter model) using GraphPad Prism v10.

A panel of peptide variants based on the native epitope (LLWNGPMAV) was synthesized in-house, with systematic substitution at each position using all standard amino acids, excluding cysteine. This panel was diluted to three times the EC90 concentration determined for each TCR.

A total of 8 x 10^6^ Jurkat-NFAT-CD8 cells, transfected with TCRs, were prepared using the Neon Transfection System 100 μL Kit (4 × 100 μL tips). Six white 96-well plates (Costar) were used for co-culture of the TCR-transfected Jurkat-NFAT-CD8 cells and T2 cells with the panel of peptide variants, in duplicate (as described above). Each plate included wells containing the native peptide and medium only as controls. After overnight incubation at 37 °C, luminescence was quantified as previously described.

For each plate, luminescence responses were normalized by setting the response to the native peptide as 100% and the response to medium only as 0% using GraphPad Prism v10.

### X-ray crystallography

Crystals were obtained by vapor diffusion using sitting-drop crystallization, Equal volumes (0.2 μL + 0.2 μL) of protein–peptide complex and reservoir solution were dispensed using a Mosquito robot (STP Labtech). Crystallization drops were incubated at 18 °C for several weeks until crystals appeared. C0-crystals of the complex with peptide YF (sequence: LLWNGPMAV) were obtained in 0.005 M cadmium chloride hemi(pentahydrate), 0.1 M Tris pH 8.5, and 20% w/v PEG 4000. Co-crystals of the S0 variant (sequence: SLLWNGPMAV) were obtained in 0.005 M nickel(II) chloride hexahydrate, 0.1 M Tris pH 7.5, and 20% w/v PEG 4000, while co-crystals of the S10 variant (sequence: LLWNGPMAVS) formed in 0.2 M ammonium nitrate, 0.1 M HEPES pH 7.5, and 20% v/v PEG Smear Broad. Crystals were cryoprotected by briefly soaking in mother liquor supplemented with 25% (v/v) glycerol and flash-frozen in liquid nitrogen. X-ray diffraction data were collected at beamline ID30B/ID30A1 of the European Synchrotron Radiation Facility (ESRF, Grenoble, France) and processed using the Autoprocess pipeline (Global Phasing) ^49–52^.

Structures were solved by molecular replacement using Phaser-MR ^53^ and PDB entry 5N6B as template model ^25^. Manual model building and structure refinement were carried out in Phenix Suite using coot software and phenix-refine, respectively. After validation, the models were deposited in the Protein Data Bank under accession numbers PDB:9SL0 (YF), PDB:9SKP (S0), PDB:9SKO (S10). Data collection and refinement statistics are summarized (Table S11). Structural representations were prepared using PyMOL (https://pymol.org).

### Predicting cross-reactivity with MixTCRcross

MixTCRcross predicts cross-reactivity between two epitopes (E1 and E2) restricted to their respective MHC (MHC1 and MHC2) by comparing the sequences of E1 and E2, the similarity between the TCR interfaces of the two MHC (positions 65, 68, 69, 72, 73, 75, 76, 146, 150, 151, 154, 157, and 158, numbering following X-ray structures), and the binding affinity of E2 to MHC2 (i.e., B2, computed as %rank of MixMHCpred ^54^). The three cross-reactivity levels (*CR*) are defined as:

-0 for at least one variant at non-anchor P3-P5, for MHCs with different TCR interfaces, or for B2 > 2, otherwise:

-1 for at least one variant at non-anchor P6-P7.

-2 for variants only at other positions.

The final score combines CR and the %rank of E2 (i.e., B2) as: 𝑆 = 𝐶𝑅 –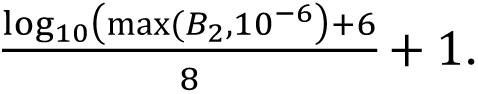 This score goes from 0 (lowest risk of cross-reactivity) to almost 3 (highest risk of cross-reactivity).

### Analysis of BATCAVE data

All data for 9-mer epitopes were downloaded from the BATCAVE database ^31^. For each TCR – peptide variant pair, activity scores were normalized by the activity of the index peptide. To benchmark how predictions of MixTCRcross can generalize to other epitopes and compare with those of BATMAN ^31^ and CrossDome ^32^ in the context of unseen epitopes, BATMAN was retrained for each index peptide on all BATCAVE data except activity measurements with variants of the index peptide (i.e., leave-one-epitope-out cross-validation), using the script kindly provided at https://github.com/meyer-lab-cshl/BATMAN-paper. CrossDome was used in its published version, which supports only a subset of the epitopes in the BATCAVE database. For AUC calculations, activity were binned in two categories based on a threshold of 10% with respect to the activity of the index peptides.

### Prediction of MAGE-A3 cross-reactive self-peptides

All human proteins were retrieved from UniProt and chopped into all possible 9-mers (11’292’067 peptides in total). MixTCRcross scores were computed for all these peptides using as input peptide (E1) the MAGE-A3 epitope (`EVDPIGHLY`), and with the HLA-A*01:01 allele.

### Analysis of the contact with TCRs mediated by peptide residues

The number of interactions with the TCRs mediated by each peptide residue was computed across all X-ray structures of human native TCRs in complex with a 9-mer peptides, based on a threshold of 5Å. Peptide positions were grouped as non-anchor P3-P5, non-anchor P6-P7 and other positions. The average number of interactions was retrieved for each set of positions and each X-ray structure (see Table S12). Average values were taken across structures with the same epitope.

### Analysis of negative data used in IMMREP23

Negative data in the IMMREP23 competition were collected for all non-A0201_GILGFVFTL epitopes. The average TSP (mean + standard deviation across all non-A0201_GILGFVFTL) was derived and compared with the TSP of TCRs recognizing A0201_GILGFVFTL.

### Interpretation of machine learning TCR-epitope interaction predictions

To interpret machine learning predictions of TCR-epitope recognition, we first generated a repertoire of 10^6^ TCRs with exactly the same VJ distribution as our baseline model for both chains. This dataset comprises both TCRs from paired TCR repertoires as well as randomly paired TCRs from single-chain repertoires. We then applied TEMPO, NetTCR, MixTCRpred and TULIP on these data with three epitopes: A0201_LLWNGPMAV, A0201_GILGFVFTL and B1302_LLWNGPMAV. The first two epitopes have large amount of training data, while the last one corresponds to a YF variant considered in this study and was used as an example of an unseen epitope (hence MixTCRpred and TEMPO were not applied with it). The top 0.1% predicted TCRs were used to display the predicted TSPs.

### AlphaFold3-based predictions of TSPs

The interactions between the YF epitope and TCRs from the ex-vivo TCRαβ repertoire from donor LAU5013 determined in this study (i.e., 6624 unique paired TCRαβs, see Table S13) were predicted using AF3 ^16^. Five inferences, using five different seeds, were performed for each TCR. The final score was defined as the sum of interchain Predicted Alignment Error (PAE) between the epitope (i.e., peptide + MHC) and the TCR (i.e., TCRα+ TCRβ). The top 1% best scoring TCRs were used to display the TSP predicted by AF3.

## Data and code availability

### Data availability

New X-ray structure were deposited in the Protein Data Bank under accession numbers PDB: 9SL0 (YF), PDB:9SKP (S0), PDB:9SKO (S10).

The raw sequencing data (fastq files) of the input and output phage-display libraries were deposited at European Nucleotide Archive (ENA) under the project number PRJEB95024. All other data are available in the supplementary tables.

### Code availability

-The TCR-epitope motif-based interaction predictor (TEMPO) is available at: https://github.com/GfellerLab/TEMPO. This tool includes all epitopes considered in this work (including some with only single-chain data that were not included in the cross-validation analysis), as well as all the YF variants.

-The cross-reactivity predictor (MixTCRcross) is available at: https://github.com/GfellerLab/MixTCRcross

-TSPs for all epitopes analyzed in this work can be interactively explored at the TCR Motif Atlas: https://tcrmotifatlas.unil.ch.

All tools are freely available for academic research. Licensing information for commercial users are included with each tool.

## Supporting information

TablsS11

TablsS12

## Acknowledgments

Library preparation and single-cell TCR sequencing was performed at the Lausanne Genomic Technologies Facility, University of Lausanne, Switzerland (https://wp.unil.ch/gtf/). We acknowledge the European Synchrotron Radiation Facility (ESRF) for provision of synchrotron radiation facilities under proposal number MX2644 and we would like to thank the beamline scientists for assistance and support in using beamline MASSIF-1 and ID30B. This project has received funding from the SNF Sinergia program (CRSII5_193749) to D.G, V.Z, A.H and G.C, the SNF Project Grant (231333) to D.G. and Y.L. We are thankful to Nathalie Rufer and Julien Schmidt for practical help and advice along the project.

## Contributions

D.G. designed the study. Y.L., A.M., A-C.T., R.G., R.L., P.G., K.L. & A.L. performed experiments. Y.L., G.C., D.M., M.P., V.Z. & D.G. analyzed the data. Y.L., G.C., D.T., J.R. & D.G., developed and implemented the computational and web-based tools, F.P., S.D., P.B., V.Z., A.H, D.G, supervised the work. M.H., D.S., P.G., J.S., A.H. provided reagents, Y.L and D.G. made the Figures. D.G wrote the manuscript with contribution from all co-authors.

