## Supplementary Figures for "Key determinants of T cell epitope recognition revealed by TCR specificity profiles"

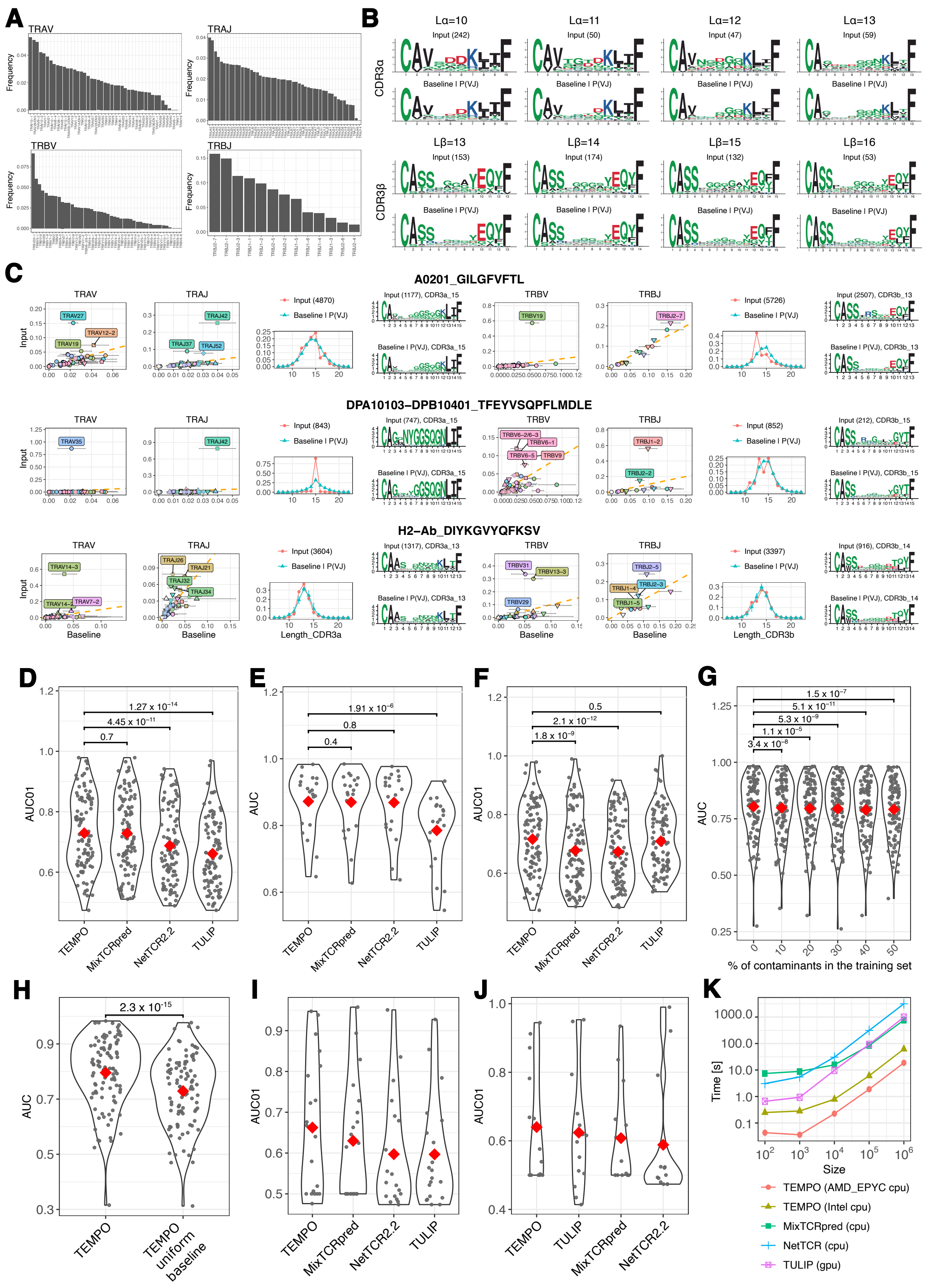

**Figure S1**. **A:** V and J gene usage in baseline TCR repertoires (average over all studies in Table S1). **B:** Comparisons of CDR3 motifs of different lengths in YF specific TCRs with the CDR3 motifs of baseline TCR repertoires with the same V/J usage (see https://tcrmotifatlas.unil.ch for other examples). **C:** TSPs for A0201_GILGFVFTL (class I Influenza epitope in human), DPA10103–DPB10401_TFEYVSQPFLMDLE (class II COVID-19 epitope in human) and H2–Ab_DIYKGVYQFKSV (class II LCMV epitope in mouse). See other examples in https://tcrmotifatlas.unil.ch. **D:** AUC01 obtained in the cross-validation for different tools trained on the same data. **E:** AUC values obtained in the cross-validation for different tools trained on the same data for epitopes with at least 200 paired TCRs. **F:** AUC01 obtained in the cross-validation for different tools trained on the same data when using as negatives in the test set TCRs found to interact with other epitopes (the so-called 'swapped negatives'). **G**: TEMPO AUC values obtained with different fractions of noise in the training data. **H:** Comparison of AUC values when using the TEMPO default baseline or a uniform baseline. **I**: AUC01 values obtained on the IMMREP23 data using the published version of different tools. **J:** AUC01 values obtained on the ePytope-TCR data using the published version of different tools. **K:** Comparison of running time as a function of the number of TCRs to predict for TEMPO and other tools.

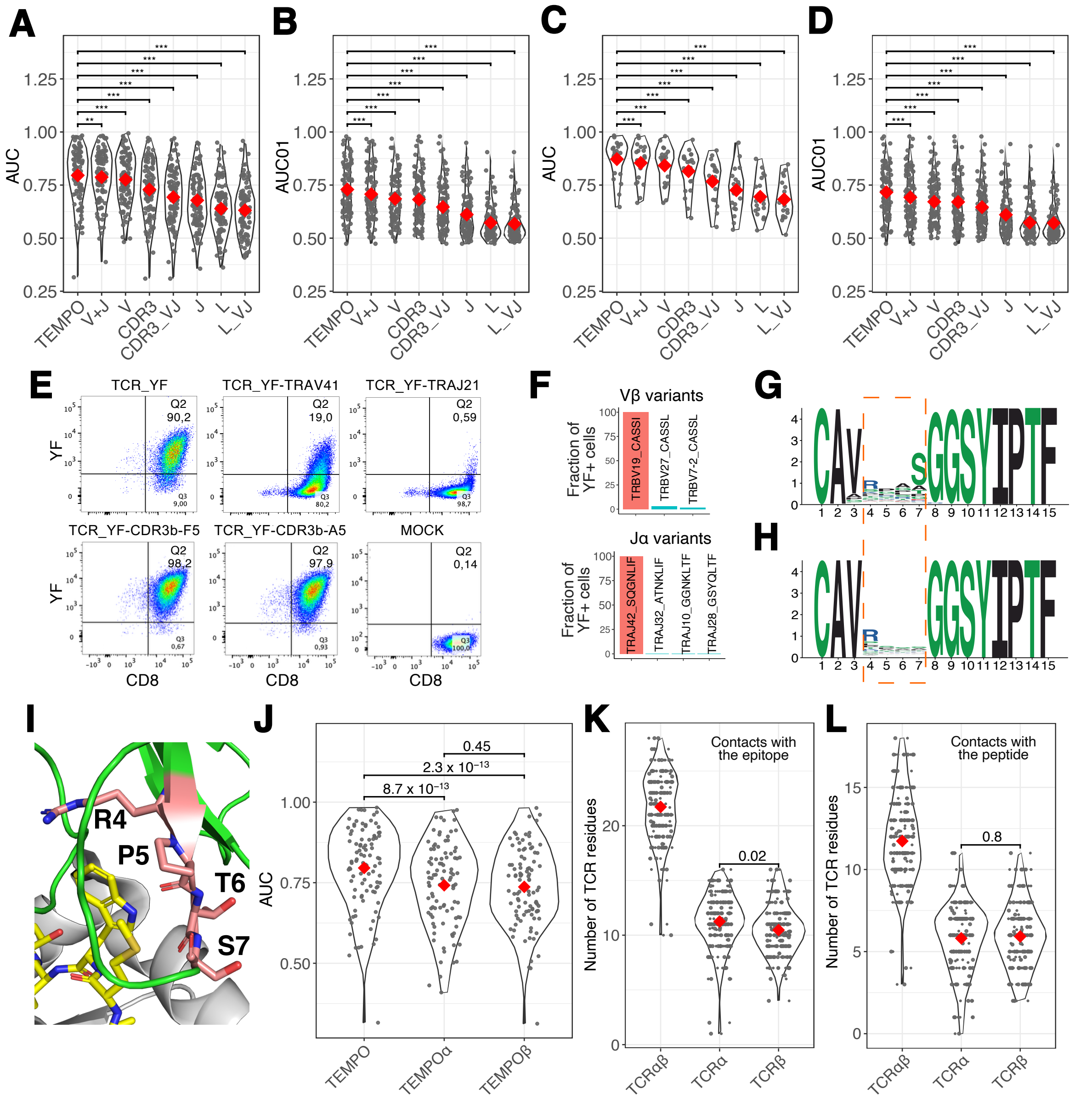


**Figure S2: A**: AUC for the cross-validation analysis with TEMPO considering different features for both chains. “CDR3_VJ” stands for the model based on CDR3 motifs ($P(CDR3|L)$) with the $Q_{P(V,J|L)}(CDR3|L)$ normalization. “L_VJ” stands for the model based on CDR3 motifs ($P(L)$) with the $Q_{P(V,J)}(L)$ normalizxation. “TEMPO” corresponds to the V+J+L_VJ+CDR3_VJ model. Chain indication is omitted for clarity. **B**: AUC01 for the cross-validation analysis with TEMPO considering different features. **C**: AUC for the cross-validation analysis when restricting to epitopes with at least 200 paired TCRs. **D**: AUC01 for the cross-validation analysis with swapped negatives in the test set. **E:** Representative FACS plots for measuring epitope recognition by transfecting into Jurkat cells different variants of TCR_YF (TRAV12-2, TRAJ30, CAVGDDKIIF, TRBV28, TRBJ2-7, CASTPQTAYEQYF)

and staining them with the YF multimer (see Table S4). **F:** Results of the staining with A0201_GILGFVFTL of Jurkat cells transfected with diverse Vβ and Jβ variants of TCR_GIL (TRAV27, TRAJ42, CAGGGSQGNLIF, TRBV19, TRBJ2-7, CASSIRSSYEQYF). **G:** CDR3α motif of baseline TCR repertoire with TRAV21, TRAJ6 and CDR3 length of 15 (i.e., $Q(CDR3_{\alpha}|\mathrm{TRAV21}, \mathrm{TRAJ6}, L_{\alpha} = 15)$). **H:** CDR3α motifs obtained in the output phage library before running MoDec. **I:** X-ray structure of the template TCR used in the phage display screen in complex with NY-ESO-1 (PDB: 2BNR). Residues at the four positions diversified in the phage display screen (R4, P5, T6, S7) are shown in orange, the peptide (SLLMWITQC) is shown in yellow and the MHC (HLA-A*02:01) is shown in grey. **J:** AUC values obtained in the cross-validation when training and testing TEMPO on both chains, only on alpha chains (TEMPOα) or only on beta chains (TEMPOβ). **K**: Number of TCRα and TCRβ residues in contact with the epitope (i.e., peptide+MHC) in experimental X-ray structures. **L**: Number of TCRα and TCRβ residues in contact with the peptide in experimental X-ray structures. In panels A-D, *** indicate P-values smaller than 0.0005 and ** indicate P-values smaller than 0.005. In panels K-L, P-values are shown between statistically independent variables.

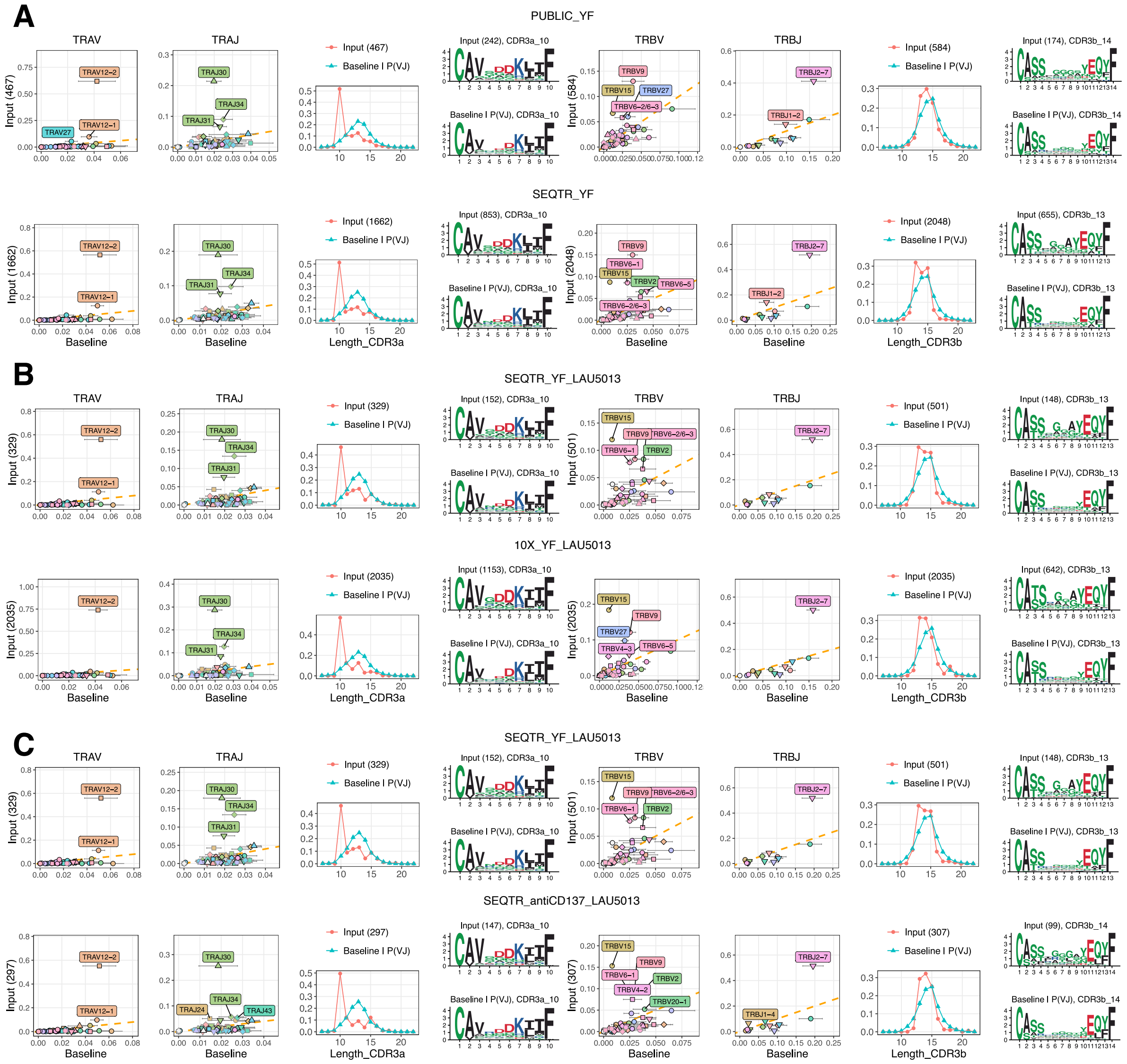


**Figure S3: A**: Comparison of the YF TSP obtained with publicly available data and the SEQTR data for the donors considered in this work (including the SEQTR specific baseline). **B**: Comparison of the YF TSP obtained with the SEQTR or 10X single-cell TCR-sequencing pipelines for donor LAU5013. **C**: Comparison of YF TSP obtained post-stimulation by sorting CD8+ cells with the YF multimer or an antibody recognizing the CD137 activation marker.

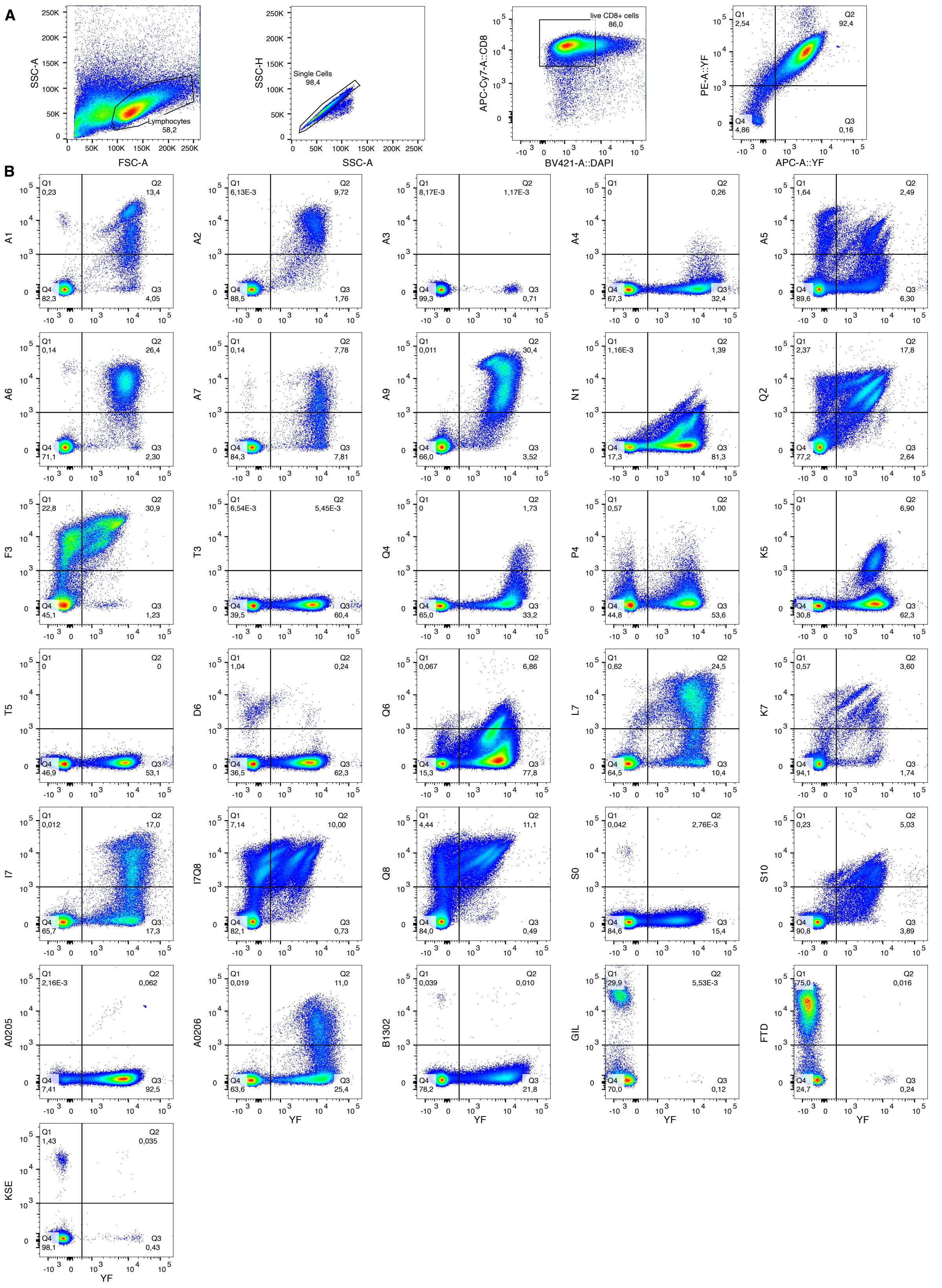


**Figure S4: A:** Gating strategy used for the cross-reactivity analyses. **B:** Cross-reactivity FACS plots for epitopes listed in Table 1, with YF multimers in APC and YF variant multimers in PE.

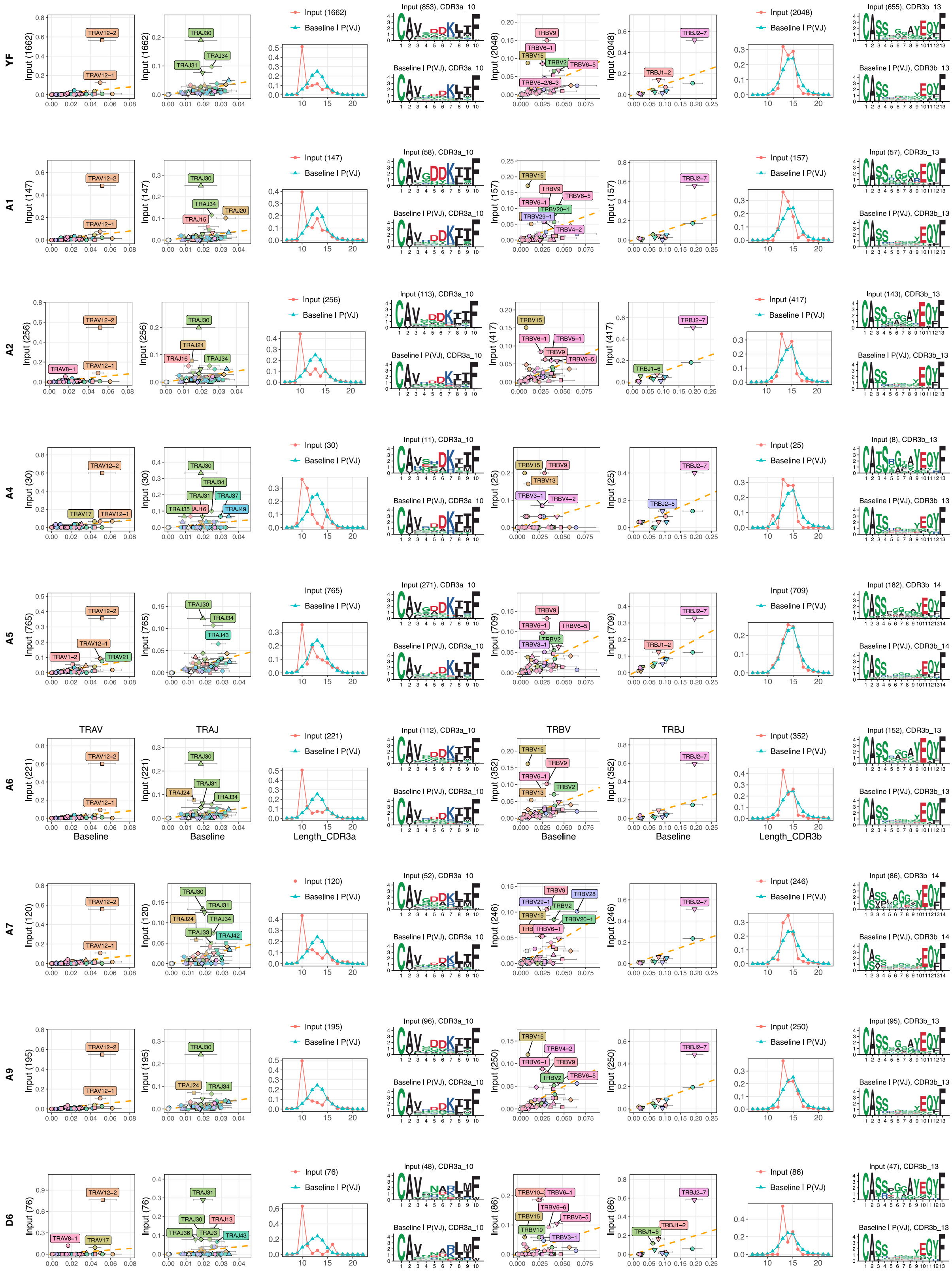

YF
A1
A2
A4
A5
A6
A7
A9
D6
TRAV
TRAJ
TRBV
TRBJ
Baseline
Length_CDR3a
Length_CDR3b
Input (1662)
Baseline I P(VJ)
Input (853), CDR3a_10
Baseline I P(VJ), CDR3a_10
Input (2048)
Input (655), CDR3b_13
Baseline I P(VJ), CDR3b_13
TRAV12−2
TRAV12−1
TRAJ30
TRAJ34
TRAJ31
TRBV9
TRBV6−1
TRBV15
TRBV2
TRBV6−5
TRBJ2−7
TRBJ1−2
Input (147)
Input (58), CDR3a_10
Input (157)
Input (57), CDR3b_13
Input (256)
Input (113), CDR3a_10
Input (417)
Input (143), CDR3b_13
Input (30)
Input (11), CDR3a_10
Input (25)
Input (8), CDR3b_13
Input (765)
Input (271), CDR3a_10
Input (709)
Input (182), CDR3b_14
Baseline I P(VJ), CDR3b_14
Input (221)
Input (112), CDR3a_10
Input (352)
Input (152), CDR3b_13
Input (120)
Input (52), CDR3a_10
Input (246)
Input (86), CDR3b_14
Input (195)
Input (96), CDR3a_10
Input (250)
Input (95), CDR3b_13
Input (76)
Input (48), CDR3a_10
Input (86)
Input (47), CDR3b_13

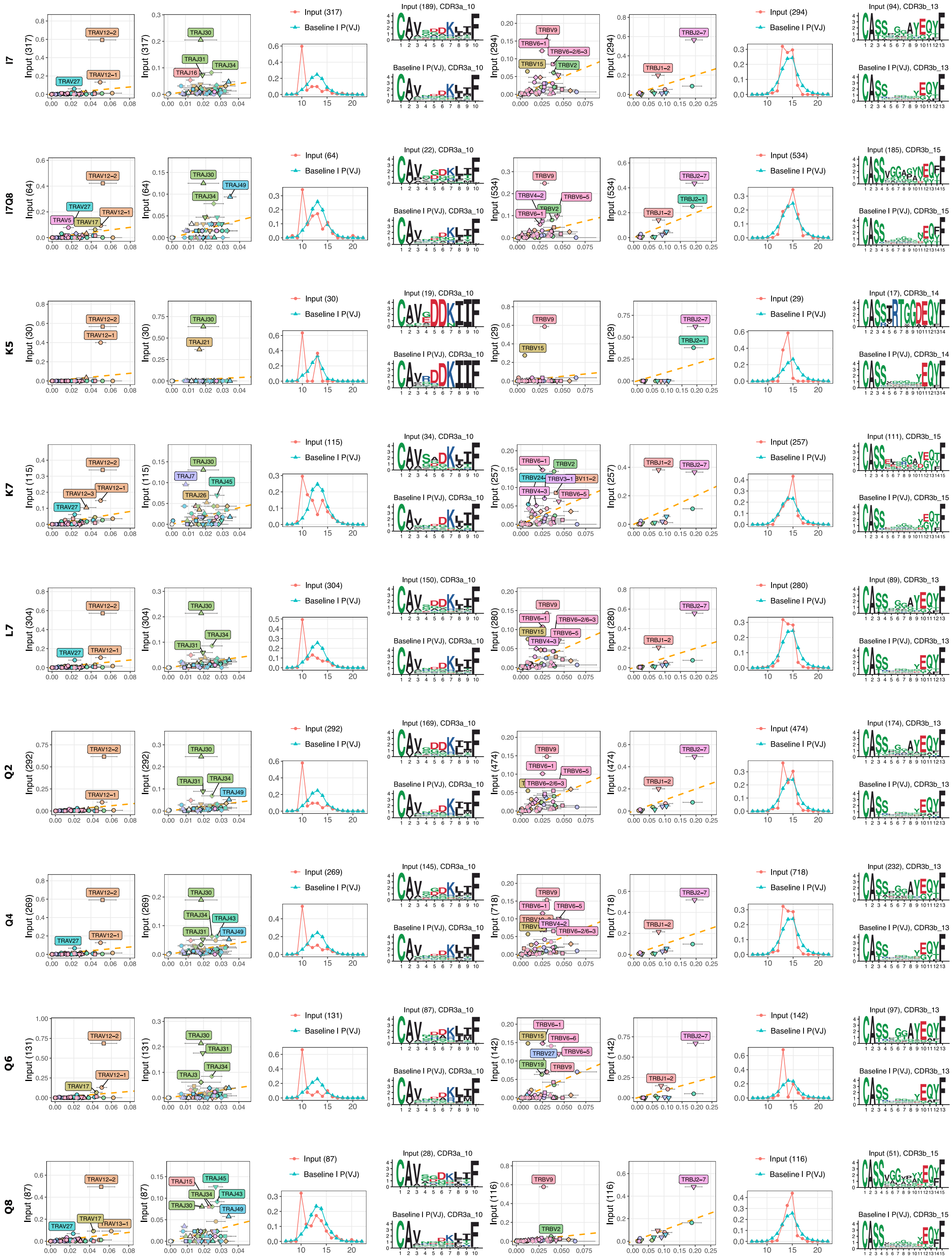

I7
Input (317)
Baseline I P(VJ)
Input (189), CDR3a_10
Baseline I P(VJ), CDR3a_10
Input (294)
Input (94), CDR3b_13
Baseline I P(VJ), CDR3b_13
I7Q8
Input (64)
Input (22), CDR3a_10
Input (534)
Input (185), CDR3b_15
Baseline I P(VJ), CDR3b_15
K5
Input (30)
Input (19), CDR3a_10
Input (29)
Input (17), CDR3b_14
Baseline I P(VJ), CDR3b_14
K7
Input (115)
Input (34), CDR3a_10
Input (257)
Input (111), CDR3b_15
L7
Input (304)
Input (150), CDR3a_10
Input (280)
Input (89), CDR3b_13
Q2
Input (292)
Input (169), CDR3a_10
Input (474)
Input (174), CDR3b_13
Q4
Input (269)
Input (145), CDR3a_10
Input (718)
Input (232), CDR3b_13
Q6
Input (131)
Input (87), CDR3a_10
Input (142)
Input (97), CDR3b_13
Q8
Input (87)
Input (28), CDR3a_10
Input (116)
Input (51), CDR3b_15

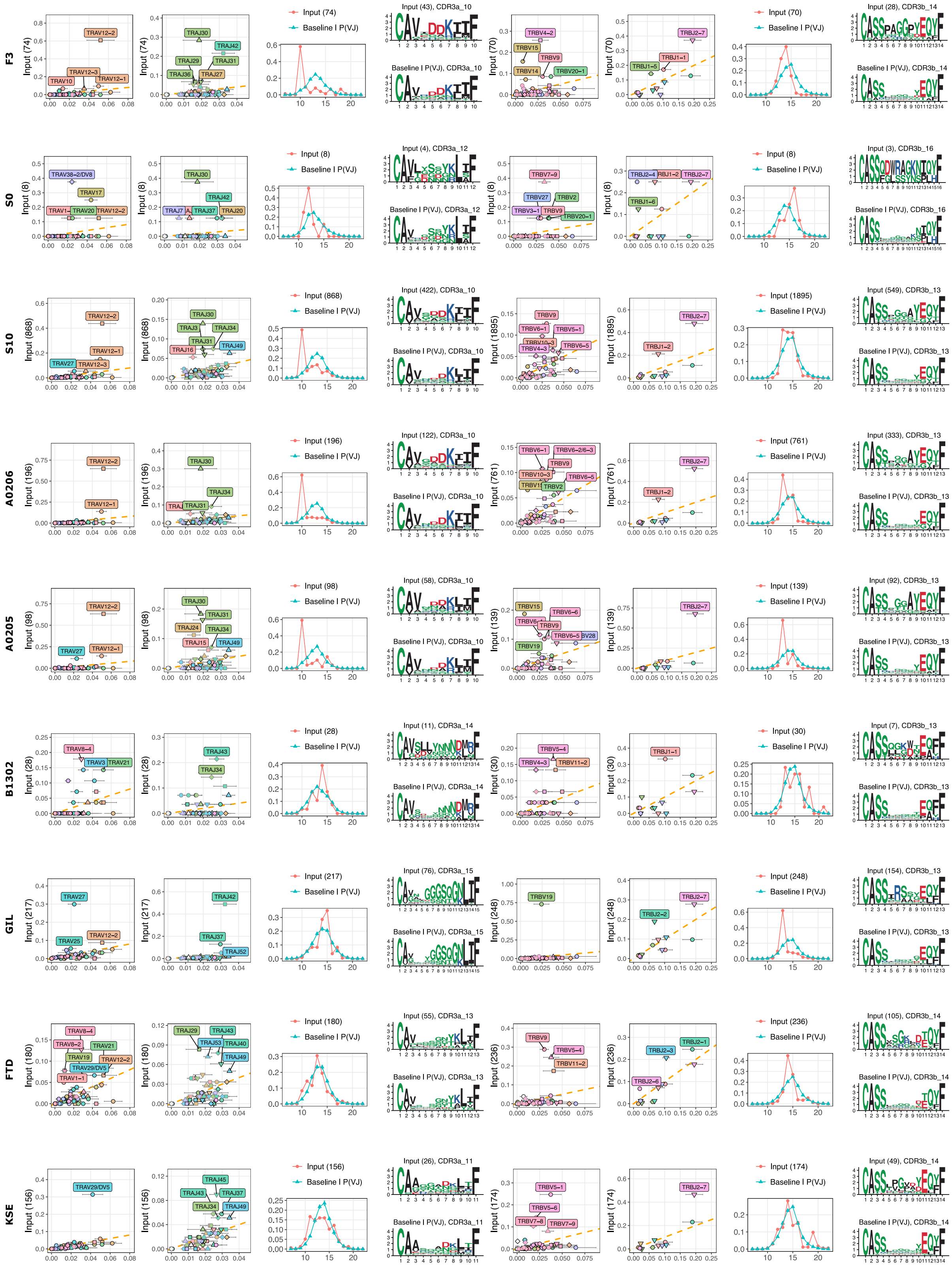


**Figure S5:** TSPs for all YF variants for which we could obtain multimer positive CD8 T cells (see also https://tcrmotifatlas.unil.ch/browse_epitopes_YF).

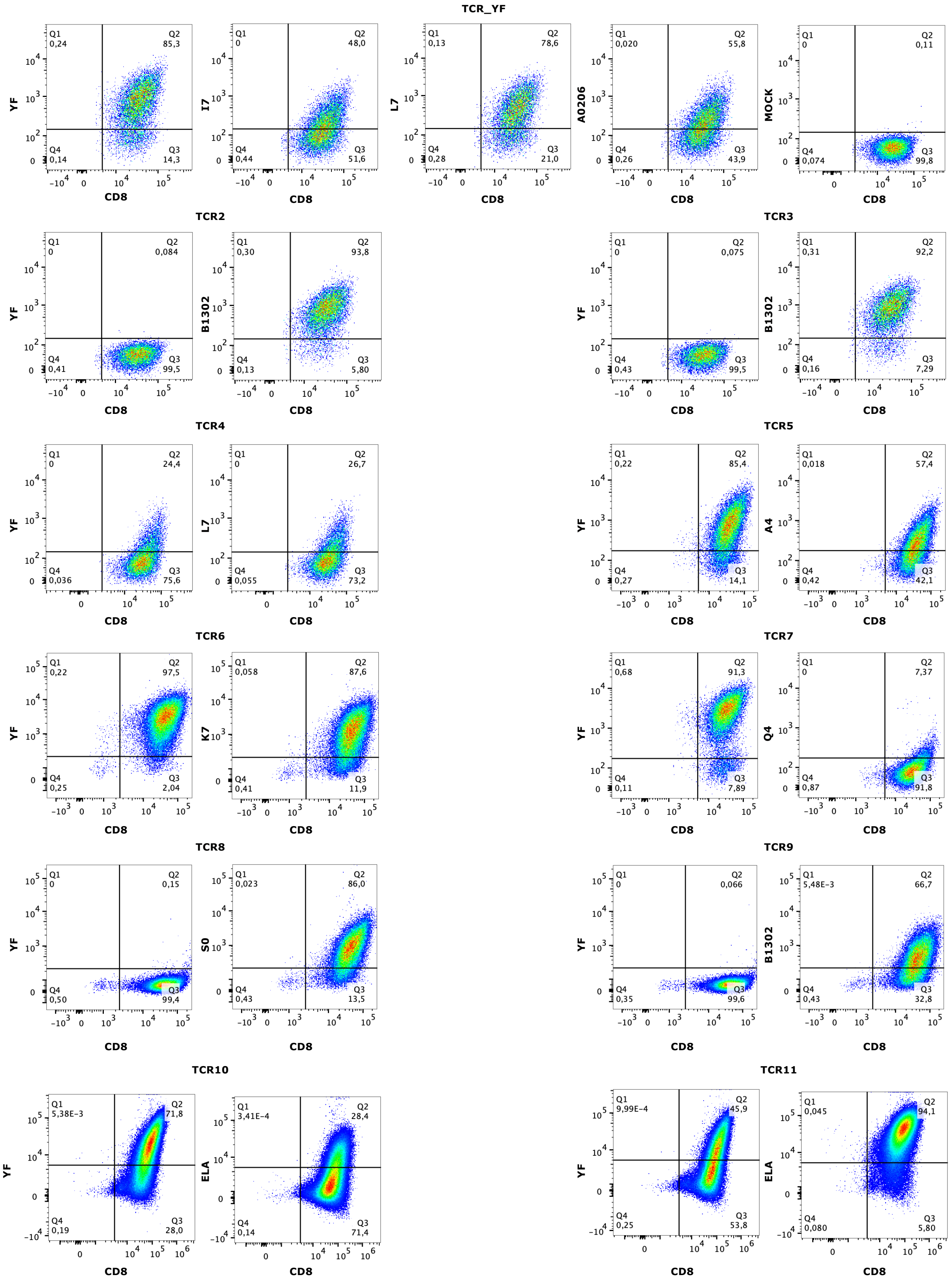
TCR_YF
YF
I7
L7
A0206
MOCK
CD8
Q1 0,24 Q2 85,3 Q4 0,14 Q3 14,3
Q1 0 Q2 48,0 Q4 0,44 Q3 51,6
Q1 0,13 Q2 78,6 Q4 0,28 Q3 21,0
Q1 0,020 Q2 55,8 Q4 0,26 Q3 43,9
Q1 0 Q2 0,11 Q4 0,074 Q3 99,8
TCR2
B1302
Q1 0 Q2 0,084 Q4 0,41 Q3 99,5
Q1 0,30 Q2 93,8 Q4 0,13 Q3 5,80
TCR3
Q1 0 Q2 0,075 Q4 0,43 Q3 99,5
Q1 0,31 Q2 92,2 Q4 0,16 Q3 7,29
TCR4
Q1 0 Q2 24,4 Q4 0,036 Q3 75,6
Q1 0 Q2 26,7 Q4 0,055 Q3 73,2
TCR5
A4
Q1 0,22 Q2 85,4 Q4 0,27 Q3 14,1
Q1 0,018 Q2 57,4 Q4 0,42 Q3 42,1
TCR6
K7
Q1 0,22 Q2 97,5 Q4 0,25 Q3 2,04
Q1 0,058 Q2 87,6 Q4 0,41 Q3 11,9
TCR7
Q4
Q1 0,68 Q2 91,3 Q4 0,11 Q3 7,89
Q1 0 Q2 7,37 Q4 0,87 Q3 91,8
TCR8
S0
Q1 0 Q2 0,15 Q4 0,50 Q3 99,4
Q1 0,023 Q2 86,0 Q4 0,43 Q3 13,5
TCR9
Q1 0 Q2 0,066 Q4 0,35 Q3 99,6
Q1 5,48E-3 Q2 66,7 Q4 0,43 Q3 32,8
TCR10
ELA
Q1 5,38E-3 Q2 71,8 Q4 0,19 Q3 28,0
Q1 3,41E-4 Q2 28,4 Q4 0,14 Q3 71,4
TCR11
Q1 9,99E-4 Q2 45,9 Q4 0,25 Q3 53,8
Q1 0,045 Q2 94,1 Q4 0,080 Q3 5,80

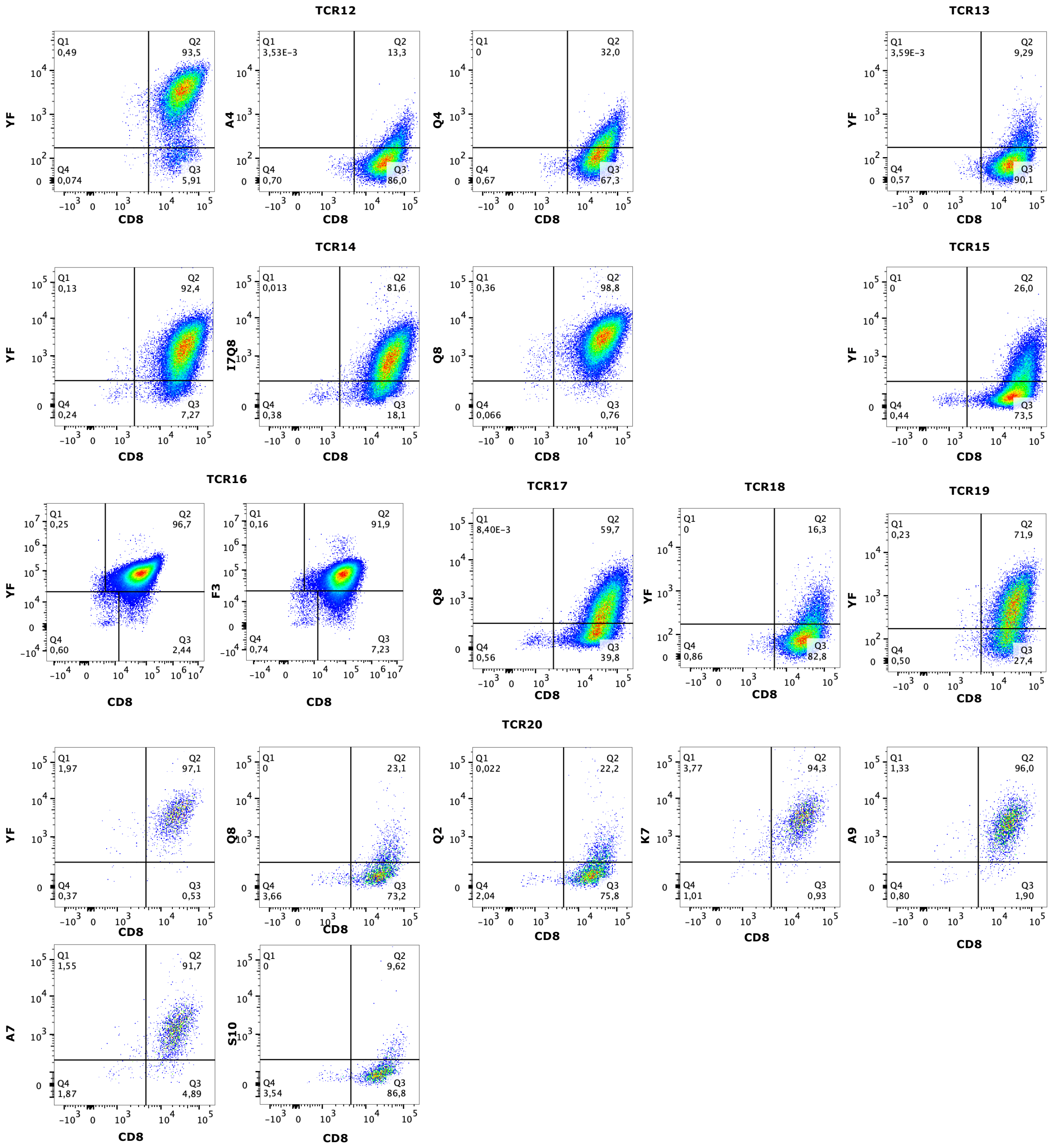


**Figure S6:** *In vitro* validation of the binding of specific TCRs observed by TCR sequencing for different variants of the YF epitope (see Table S9).

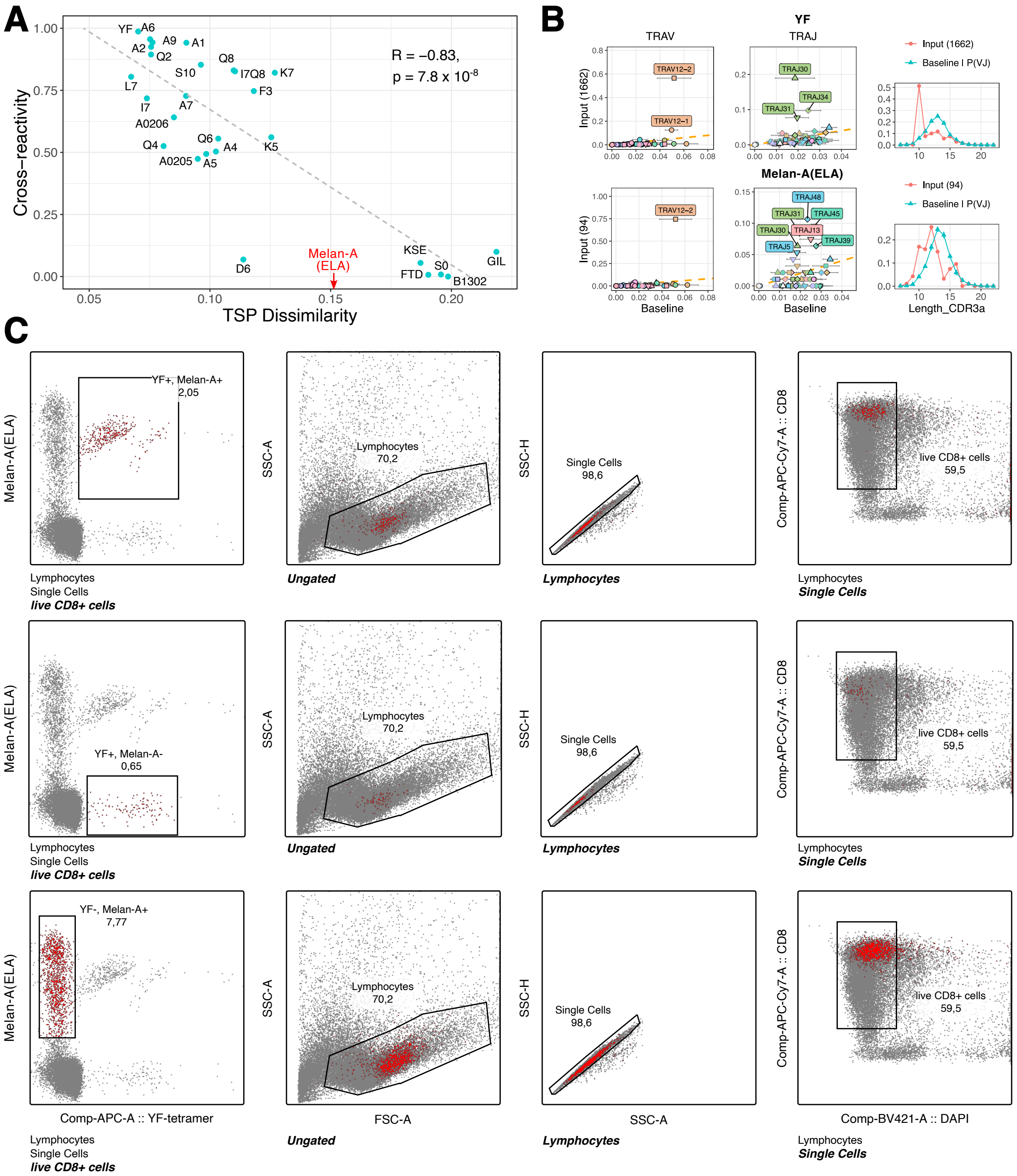


**Figure S7: A:** Correlation between cross-reactivity and TSP dissimilarity when including CDR3_VJ contribution in the TSP dissimilarity measure. **B:** Comparison between the YF and the Melan-A (A0201_ELAGIGILTV) TSPs. **C:** Gating strategy for the cross-reactivity analysis between the YF and Melan-A (A0201_ELAGIGILTV) epitopes.

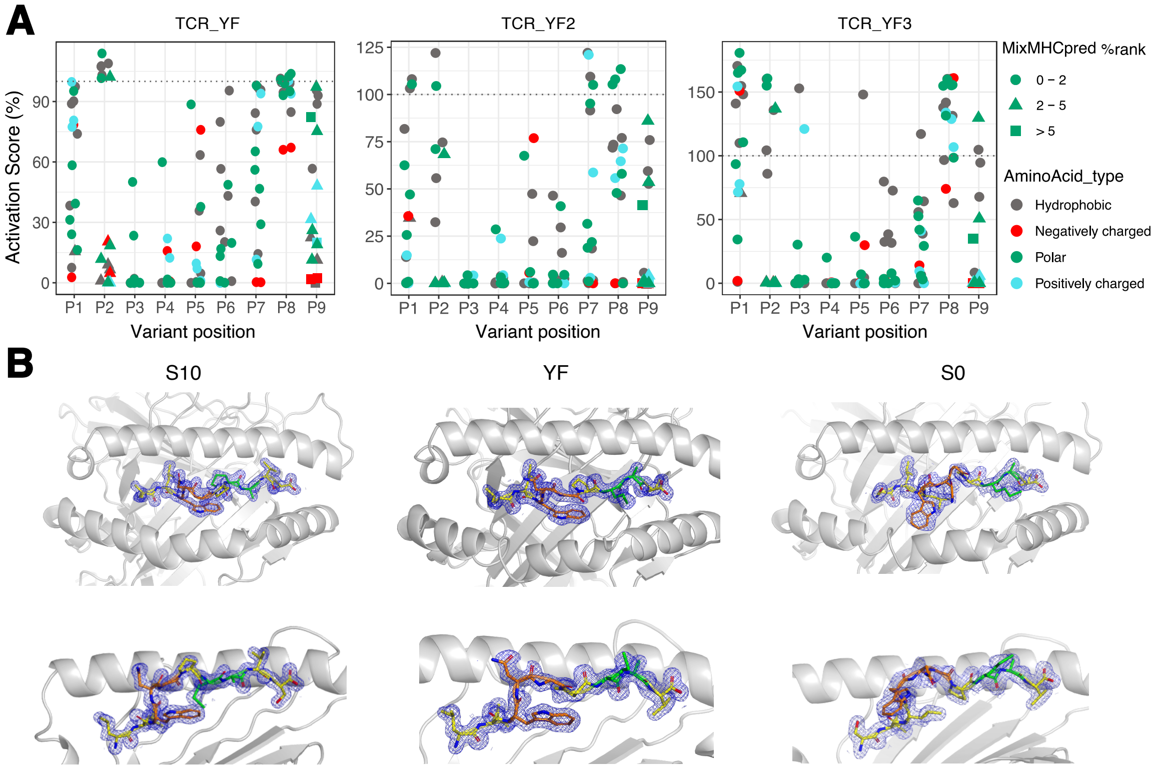


**Figure S8: A**: Results of the X-scan for three YF specific TCRs (TCR_YF: TRAV12-2, CAVGDDKIIF, TRAJ30, TRBV28, CASTPQTAYEQYF, TRBJ2-7; TCR_YF2: TRAV12-2, CAVNPDKIIF, TRAJ30, TRBV4-2, CASSQEDRGPEKLFF, TRBJ1-4; TCR_YF3: TRAV12-2, CAAGDDKIIF, TRAJ30, TRBV29-1, CSVATSGGSNEQFF, TRBJ2-1). **B:** Electron density of X-ray structure for the S10 variant, the YF and the S0 variants. For each epitope, a top view and a side view are displayed.

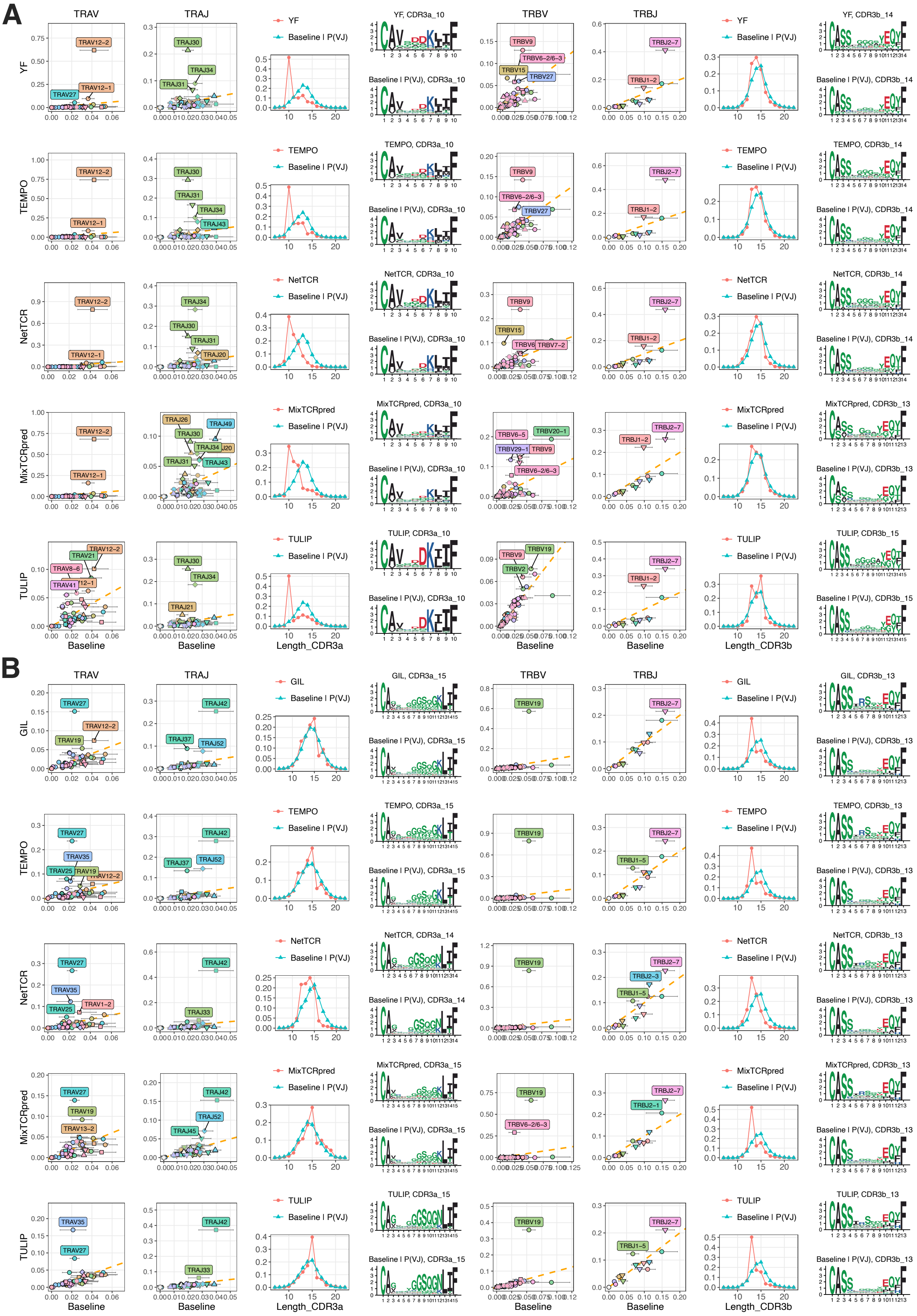
A
B
TRAV
TRAJ
TRBV
TRBJ
YF
TEMPO
NetTCR
MixTCRpred
TULIP
GIL
Baseline | P(VJ)
YF, CDR3a_10
Baseline | P(VJ), CDR3a_10
YF, CDR3b_14
Baseline | P(VJ), CDR3b_14
TEMPO, CDR3a_10
TEMPO, CDR3b_14
NetTCR, CDR3a_10
NetTCR, CDR3b_14
MixTCRpred, CDR3a_10
MixTCRpred, CDR3b_13
Baseline | P(VJ), CDR3b_13
TULIP, CDR3a_10
TULIP, CDR3b_15
Baseline | P(VJ), CDR3b_15
GIL, CDR3a_15
Baseline | P(VJ), CDR3a_15
GIL, CDR3b_13
TEMPO, CDR3a_15
TEMPO, CDR3b_13
NetTCR, CDR3a_14
Baseline | P(VJ), CDR3a_14
NetTCR, CDR3b_13
MixTCRpred, CDR3a_15
MixTCRpred, CDR3b_13
TULIP, CDR3a_15
TULIP, CDR3b_13
Baseline
Length_CDR3a
Length_CDR3b

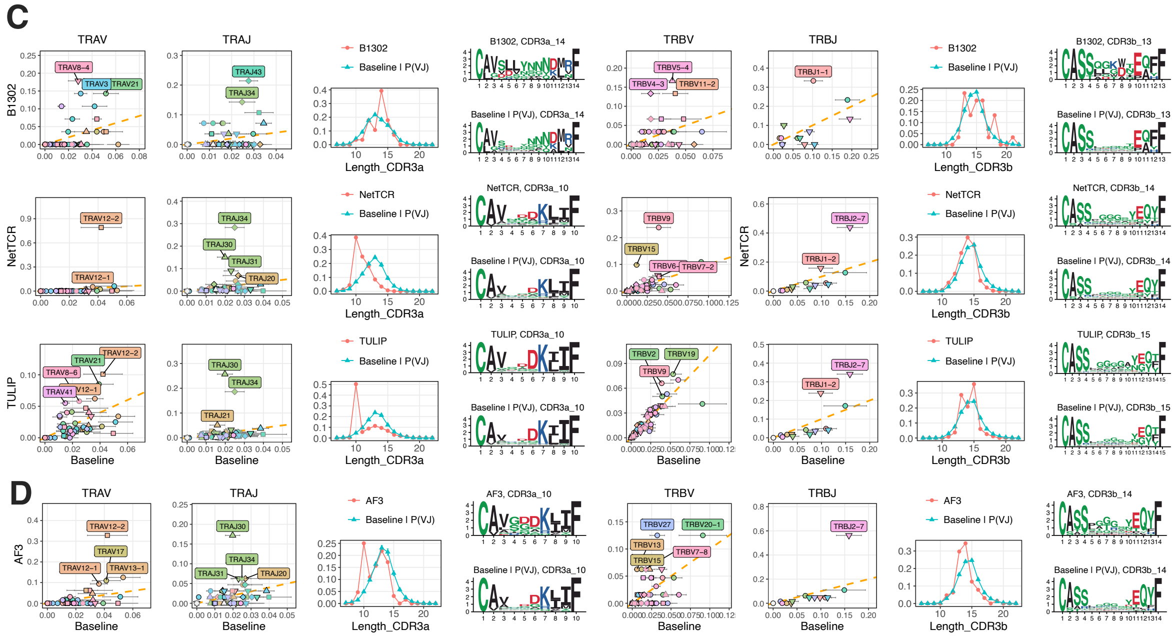


**Figure S9: A**: TSPs predicted by TEMPO, NetTCR, MixTCRpred and TULIP for the YF epitope. **B:** TSPs predicted by TEMPO, NetTCR, MixTCRpred and TULIP for the GIL epitope. **C**: TSPs predicted by NetTCR and TULIP for B1302_LLWNGPMAV. **D:** TSP predicted by AF3 for the YF epitope.

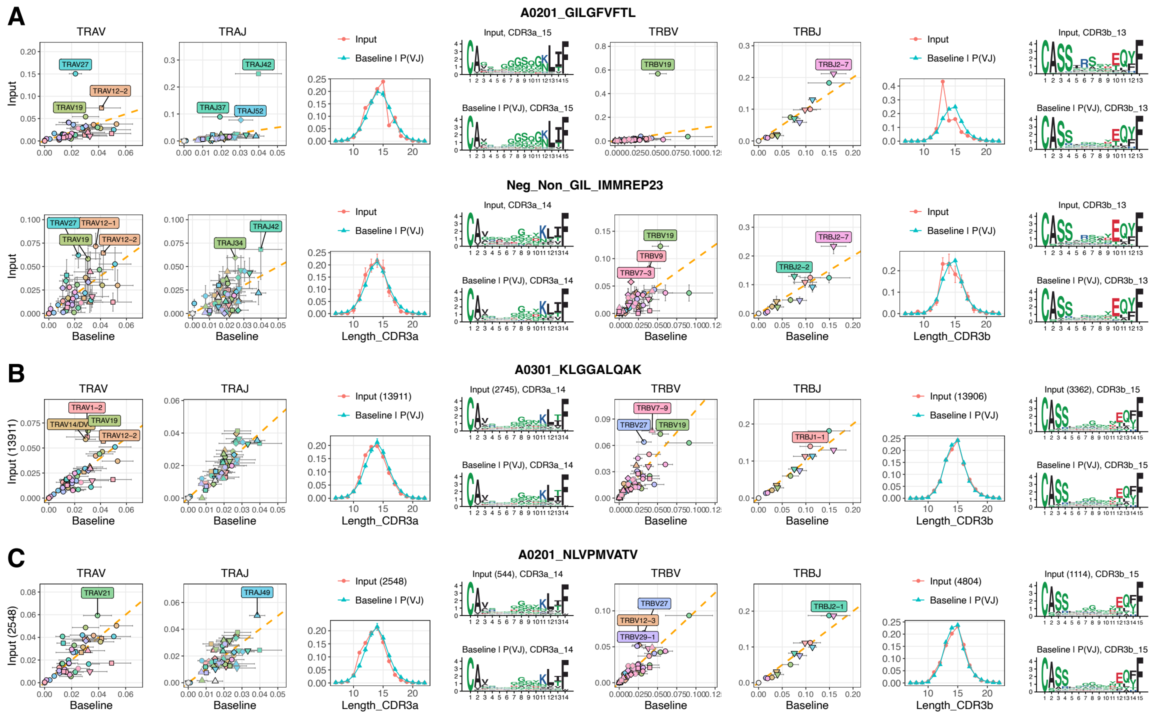


**Figure S10: A**: Comparison of the TSP for GIL and the TSP for negatives used in non-GIL epitopes in the IMMREP23 dataset (average across all non-GIL epitopes). **B**: TSP for the A0301_KLGGALQAK epitope from the 10X Genomics dataset . **C**: TSP for the A0201_NLVPMVATV epitope.

### Supplementary tables

**Table S1:** List of baseline TCR repertoire studies used to establish the baseline model (denoted with letter *Q*) in the TSPs.

**Table S2:** List of publicly available TCR – epitope interactions used in this work.

**Table S3: A**: List of TCR-epitope PDB structures with the number of contacts with the epitope (i.e., peptide + MHC) mediated by residues encoded in the V segment, the J segment or the part of the CDR3 resulting from insertions at the VJ junction (referred to as “CDR3_VJ”). Small letters in CDR1, CDR2 and CDR3 sequences indicate residues making direct contacts with the peptide. **B**: List of PDB structures with the number of contacts with the peptide mediated by residues encoded in the V segment, the J segment or the part of the CDR3 resulting from insertions at the VJ junction. Small letters in CDR1, CDR2 and CDR3 sequences indicate residues making direct contacts with the peptide.

**Table S4:** Fraction of multimer-positive Jurkat cells transfected with diverse TCRs and stained with the YF or GIL multimers.

**Table S5:** Results of the phage display experiments with a library of diversified CDR3$\alpha$ (position 4 to 7) selected against the NY-ESO-1 (A0201_SLLMWITQC) monomer. “Input” stands for the input library prior to any selection. “Output” stands for the output library after selection with NY-ESO-1 monomers.

**Table S6:** TCR sequences identified by SEQTR after sorting CD8 T cells with the YF epitope, variants or the YF epitope, unrelated epitopes or an antibody recognizing the CD137 activation marker.

**Table S7:** Results of the single-cell TCR-seq with 10X Genomics for YF-specific CD8 T cells in donor LAU5013.

**Table S8**: Cross-reactivity values for each epitope in Table 1.

**Table S9**: Fraction of multimer-positive Jurkat cells transfected with diverse TCRs and stained with the YF variants or Melan-A.

**Table S10**: Results of the X-scan assay on three TCRs recognizing the YF epitope.

**Table S11:** PDB report for the three X-ray structures (S10, YF, S0).

**Table S12:** Average number of interactions with TCR residues mediated by peptide residues at different positions (i.e., non-anchor P3-P5, non-anchor P6-P7, other positions) for X-ray structures of 9-mer epitopes in complex with human TCRs. For each epitope, values were averaged across all structures with this epitope.

**Table S13:** Results of the single-cell TCR-seq with 10X Genomics for CD8 T cells collected *ex vivo* (i.e., prior to any stimulation) from donor LAU5013. Only complete single-alpha and single-beta TCRs are reported.
