## Supplementary material for "Key determinants of T cell epitope recognition revealed by TCR specificity profiles": TablsS11

|  | **YF** | **S0** | **S10** |
| --- | --- | --- | --- |
|  | *native* | *native* | *native* |
| **Data collection** |  |  |  |
| Wavelength (Å) | 0.87313 | 0.87313 | 0.96546 |
| Space group | P 21 21 21 | P 21 21 21 | P 21 21 21 |
| *a, b, c* (Å) | 60.181, 80.201, 111.589 | 60.626, 79.959, 109.38 | 59.928, 79.197, 111.4 |
| α, β, γ (°) | 90.0, 90.0, 90.0 | 90, 90, 90 | 90, 90, 90 |
| Resolution (Å) * | 19.744 - 1.596 (1.623 – 1.596) | 19.68 - 1.894 (1.927 - 1.894) | 19.662 - 1.486 (1.512 - 1.486) |
| R_meas_* | 0.085 (1.798) | 0.154 (1.235) | 0.084 (0.892) |
| R_merge_* | 0.081 (1.730) | 0.148 (1.190) | 0.074 (0.778) |
| Mean I/σI* | 18.6 (1.6) | 13.1 (2.5) | 10.6 (1.8) |
| Completeness (%)* | 99.9 (98.9) | 100.0 (99.9) | 98.4 (99.2) |
| Multiplicity* | 13.6 (13.4) | 13.6 (13.7) | 3.9 (4.0) |
| CC1/2* | 0.999 (0.752) | 0.999 (0.721) | 0.997 (0.581) |
| **Refinement** |  |  |  |
| Resolution (Å) | 19.744 – 1.596 | 19.68 – 1.894 | 19.662 - 1.486 |
| Total reflections* | 986307 (47242) | 583525 (28580) | 340101 (16905) |
| Total unique* | 72436 (3522) | 42938 (2079) | 86353 (4273) |
| *R_work_^#^* | 0.2339 (0.2944) | 0.1859 (0.2544) | 0.1843 (0.2626) |
| *R_free_^#^* | 0.2637 (0.3286) | 0.2287 (0.3188) | 0.2041 (0.2882) |
| Number of non-hydrogen atoms | 3397 | 3497 | 3526 |
| macromolecules | 3183 | 3186 | 3173 |
| ligands | 14 | 61 | 103 |
| solvent | 200 | 250 | 250 |
| Protein residues | 382 | 383 | 383 |
| RMS deviations (bonds)^#^ | 0.006 | 0.007 | 0.005 |
| RMS deviations (angles)^#^ | 0.78 | 0.87 | 0.79 |
| Ramachandran favored (%)^#^ | 98.67 | 98.94 | 98.94 |
| Ramachandran allowed (%)^#^ | 1.33 | 1.06 | 1.06 |
| Ramachandran outliers (%)^#^ | 0.00 | 0.00 | 0.00 |
| Rotatmer outliers (%)^#^ | 0.59 | 0.30 | 0.60 |
| Clashscore^#^ | 0.62 | 6.02 | 2.36 |
| Average B-factor^#^ | 25.74 | 26.43 | 19.19 |
| macromolecules | 25.43 | 25.74 | 18.10 |
| ligands | 43.37 | 38.97 | 38.29 |
| solvent | 29.43 | 32.20 | 25.20 |
| Molprobity Score | **1.10** | **1.47** | **1.20** |
| PDB | **9SL0** | **9SKP** | **9SKO** |
| ^#^as reported by phenix.table_one and phenix.model_vs_data  *as reported by autoPROC and phenix.table_one | | |  |
